# Neural dynamics hierarchy in motor cortex and striatum across naturalistic behaviors

**DOI:** 10.1101/2024.12.17.629049

**Authors:** David Xing, Joshua Glaser, Andrew Miri

## Abstract

Mammals perform a wide range of movements actuated by diverse patterns of muscle activity. Primary motor cortex (M1) and striatum are implicated in controlling these movements, but how their activity dynamics are organized to accommodate such diversity is poorly understood. We developed a paradigm that enabled us to investigate neural dynamics across diverse motor behaviors in mice. In contrast to existing views, we found neither behavior-specific nor behavior-invariant organization in single-neuron activity, population-level covariation, and muscle activity correlation. Instead, the similarity of activity dynamics between behaviors varied differentially across behavior pairs, forming a hierarchical organization. The same hierarchical organization was shared between M1 and striatum, despite stronger muscle activity correlation in M1 and greater behavior specificity in striatum. Network modeling indicated that striatal activity is sufficient to drive hierarchical dynamics in muscle pattern-generating circuits. These hierarchical dynamics may reflect a tradeoff between behavioral specialization and generalization in motor system function.

## Introduction

Natural behavior is composed of a wide variety of motor actions. For example, a mouse may perform a series of grooming movements, switch to locomotion to approach a seed on the ground, reach for and grab the seed, freeze when a predator is detected, then dart away and climb up a tree to escape. A hallmark of the nervous system is its ability to generate the diverse and complex patterns of muscle activity that achieve these different actions. Yet it remains poorly understood how neural activity dynamics coordinated across motor system regions are organized to efficiently control such a diversity of movements. This knowledge gap impedes the development of mechanistic models of motor system operation^1,2^.

Previous work has suggested different principles that may govern this organization. One possibility is that distinct neuronal populations may be specialized for, and specifically engaged during, different motor behaviors^3–5^. Indeed, studies have identified populations in the primary motor cortex (M1) and the brainstem that are selectively active during different types of movement like walking and grooming^6^, or reaching and food handling^7–9^. Indications of behavior specificity have also been seen in activity at the population level, where the covariation of firing patterns across neurons has been observed to change between reaching and grasping^10^, co-contraction and alternation of antagonistic muscles^11^, similar movements across constrained and freely- initiated contexts^12^, and movements that differ in the degree of M1 involvement^13^.

Alternatively, the motor system could utilize a behavior-invariant organization, where neural activity encodes features of movement consistently across behaviors. Early recordings in M1 aimed to characterize the tuning of cortical neurons to particular facets of movement such as end point kinematics^14–16^, movement trajectories^17,18^, posture^19–21^, and limb kinetics^22–24^. Implicit in these studies was the goal of uncovering a stable representation of movement parameters or muscle commands^21,25^. Mor e recent studies have found that different motor tasks, like reaches in different directions^26,27^ or wrist activation that generates either movement or isometric torque^28^, share invariant patterns of population-level activity covariation. Behavior specificity and invariance can also represent two extremes on a continuum^29^; rather than adhering exclusively to either strategy, motor circuits might utilize an intermediate organization that partially leverages the advantages of each one.

However, the difficulty of studying the motor system across a broad range of behaviors has precluded assessment of the extent to which motor system activity exhibits behavior-specific or behavior-invariant organization. Motor system studies have historically focused on one or a few movement types, typically those that are cued, target-driven, highly constrained, and extensively trained^26,28^. Open field paradigms allow for the expression of more numerous and less constrained behaviors^30–33^, but these paradigms neglect much of the complexity and dexterity of the movements that animals express in natural environments^34^. Characterizing neural activity dynamics during naturalistic behaviors also requires simultaneous measurement of activity across many neurons at high temporal resolution, something that remains challenging in less constrained paradigms. Thus, the degree of similarity of neural dynamics across much of an animal’s motor repertoire remains opaque. In addition, the organization of these dynamics may also vary across motor system regions. For example, the striatum historically has been implicated in action selection^35–39^ and therefore may exhibit greater behavior specificity than spinally-projecting regions like M1, whose dynamics could reflect different functions.

To address this, we developed a new paradigm that enables the quantification of neural dynamics across diverse naturalistic behaviors in mice. These behaviors include both stereotyped, species- typical behaviors like eating and grooming, and those that challenge agility and dexterity, like climbing and walking across an irregular grid. We found that on both a single-neuron and population level, activity in forelimb M1 and striatum did not exhibit exclusively behavior-specific or behavior-invariant organization. Instead, activity dynamics during different behaviors were similar to varying degrees, with some pairs of behaviors exhibiting relatively similar dynamics and other pairs exhibiting highly dissimilar dynamics. This broad variation in neural dynamics similarity can be described using hierarchical clustering, reflecting a hierarchical organization^30,33,40,41^ in which different levels of the hierarchy represent different degrees of similarity in neural dynamics. This hierarchy was similarly organized in both M1 and striatum, despite M1 activity being more strongly correlated with muscle activity and striatal activity exhibiting greater behavior specificity. Network modeling demonstrated that observed striatal activity is able to induce graded variation across behaviors in the activity dynamics of muscle pattern-generating circuits, which constitutes a hierarchy. Collectively, our results reveal a hierarchical organization of motor system activity dynamics across behaviors, which may reflect a tradeoff between behavioral specialization and the efficiency of reusing functional elements.

## Results

### An arena paradigm for examination of diverse naturalistic motor behavior

To promote the expression of a wide range of naturalistic motor behaviors, we designed and built an arena featuring various interactable elements (Fig. 1a). These included floors with separate flat and randomly spaced, raised grid sections, walls with randomly spaced mesh handholds, and an aversive “spike pit” area to promote jumping. Mice also ate sunflower seeds that were periodically placed into the arena. To encourage continuous exploration and frequent climbing, water ports were placed at various locations around the walls, with only a single randomly chosen port active at any given time (Extended Data Fig. 1). Mice exhibited a diverse range of behaviors, including climbing, rearing, grooming, walking, eating, jumping, and stillness, which we manually categorized into 10 distinct classes using video-based annotation (Fig. 1b). We assigned time points to one of these 10 classes, or an “unlabeled” catch-all class. Climbing was the only behavior that required explicit training; all other behaviors were expressed naturally within the first session mice spent in the arena. Sessions lasted 81 ± 8 minutes (mean ± std).

**Figure 1.**
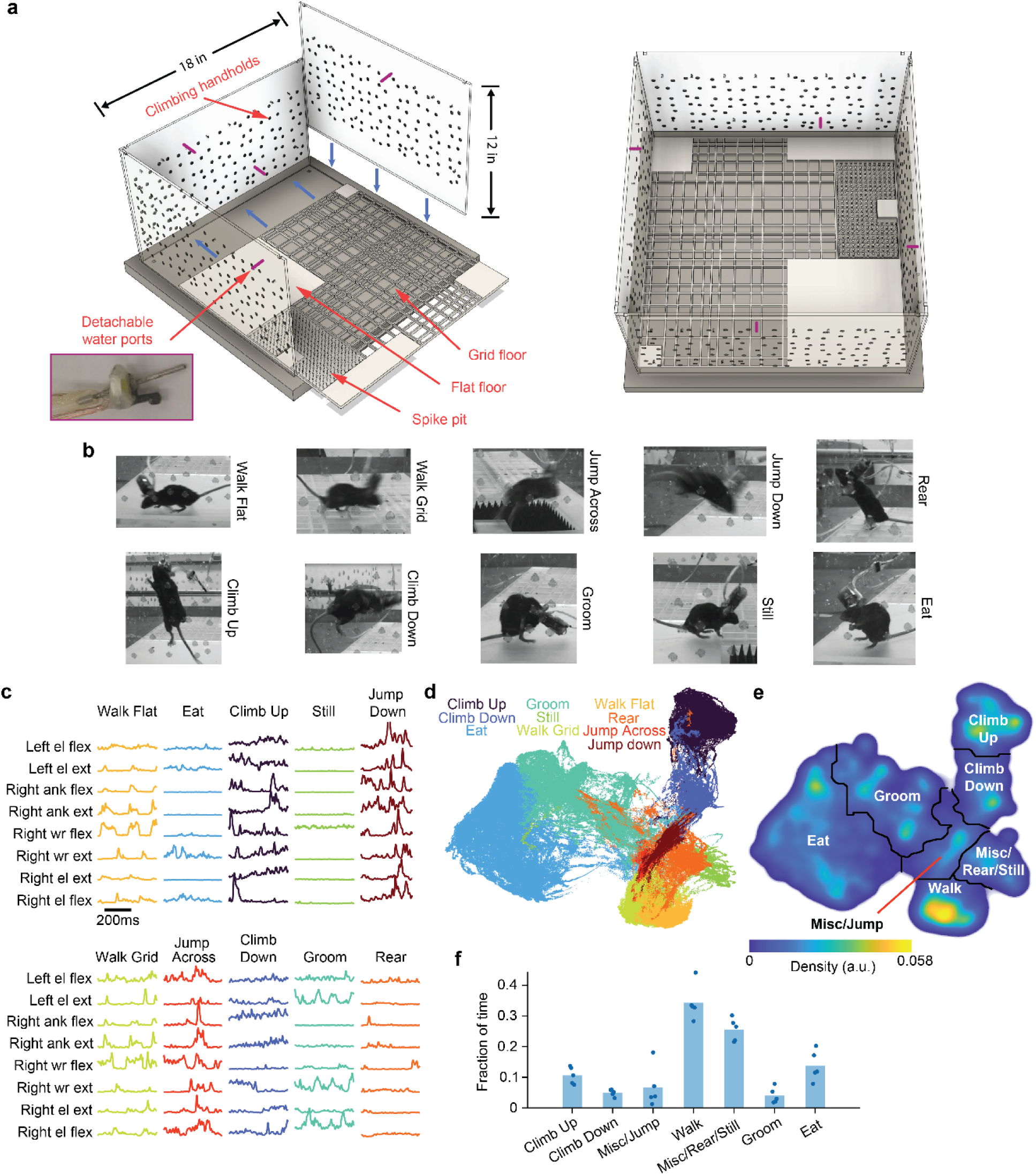
| A novel freely-behaving paradigm for interrogating naturalistic behavior. a, Left, arena design consisting of elements such as grid floors, climbing walls, and jumping regions to promote a diversity of movements. Purple lines indicate example locations for water ports. Right, top view of a fully assembled arena. **b,** Example frames for the ten different behavior classes. **c,** Example activity from eight muscles across the ten different behavior classes. Flex, flexor; ext, extensor; el, elbow; ank, ankle; wr, wrist. **d,** UMAP projection of muscle frequency features, for the time points with a manually annotated behavior class. Each point corresponds to a 1ms time bin, colored by the associated behavior class. **e,** Watershed segmentation on the density plot of the time points, followed by grouping and assignment of regions into seven behavioral regions. **f,** Mean fraction of time spent in each of the behavioral regions. Each point corresponds to an individual mouse.

To quantify motor output throughout these behaviors, we performed electromyographic (EMG) recordings from 6-8 muscles across both forelimbs and the right hindlimb^13,42^. These recordings yielded rich and diverse patterns of muscle activity such as synchronized bursts during jumping, large weight-bearing contractions during climbing, and rhythmic oscillations during walking and grooming (Fig. 1c). Using these recordings, we adapted a previously established method^43^ to obtain a 2D map of the muscle activity states for each mouse. A time-frequency transformation was applied to the EMG recordings, and frequency features at each time point were projected into two dimensions using UMAP (Extended Data Fig. 2a). Despite being agnostic to our behavior annotations, the resulting maps exhibited distinct clusters that aligned with the 10 behavior classes (Fig. 1d), similarly across mice (Extended Data Fig. 2b).

Because our manual annotation was based on fairly stringent criteria, only 25% of time points were assigned a behavior class, while the rest were assigned as unlabeled. To utilize data from all time points in our subsequent analyses, we defined behavioral regions within the UMAP-based 2D maps. We applied the watershed algorithm to the density of time points plotted across the maps, segmenting each map into seven regions (Fig. 1e). We then assigned all time points, including unlabeled points, to one of the seven behavioral regions based on their location in the UMAP projection. Many of the unlabeled time points involved small or miscellaneous movements such as quick postural adjustments, sniffing, or brief (<500 ms) pauses, behaviors that generally fell into the rearing/still or jumping regions (Fig. 1f).

For many of the observed behaviors, it is unknown whether cortex directly influences muscles through descending pathways^13^. We therefore examined whether muscle activity within each behavioral region exhibits short latency effects following rapid optogenetic inactivation of the caudal forelimb area (CFA; forelimb primary motor and somatosensory cortex). Following established methods, we applied 473 nm light pulses (50 ms duration, jittered spacing 1.0-1.3 s apart) onto the surface of CFA in transgenic mice expressing channelrhodopsin2 in vGAT+ cortical interneurons^44^ (16 sessions from three mice; Extended Data Fig. 3a). This technique has been shown to cause fast and nearly complete silencing of excitatory projection neurons in cortex^13,45,46^. Inactivation caused rapid (<35 ms) changes in contralateral forelimb muscle activity compared to controls (Extended Data Fig. 3a,b). Effects were significant in all behavioral regions at both short (35 ms) and long (100 ms) latencies, except for the misc/rear/still region (Extended Data Fig. 3c,d). Thus, our novel paradigm elicits a diverse range of motor behaviors, with most involving direct cortical influence.

### Cortical and striatal neurons exhibit a continuum of behavioral specificity

We next examined neural activity in both forelimb M1 and striatum across the different behavioral regions. Using chronically implanted Neuropixels^47^, we were able to simultaneously record from hundreds of neurons across both areas (3 sessions from 3 mice, Extended Data Fig. 4a-e). Firing rate time series for the vast majority of neurons in both regions showed a significant dependence on muscle activity, although the fraction in M1 was higher (Extended Data Fig. 4f).

To examine the relationship between each neuron’s activity and the animal’s behavior state, we calculated the mean firing rate of each neuron across points on a 1001x1001 grid over the 2D state map (Fig. 2a, Supplementary Video 1). We also computed the mean firing rate of each neuron within each of the seven behavioral regions. In a behavior-specific organization where different behaviors involve specific neuronal subpopulations, we would expect to see the majority of neurons selectively active within individual behavior regions. On the other hand, observing activity across all behavioral regions in most neurons would suggest a behavior-invariant organization. Our results did not agree with either of these predictions: in both M1 and striatum, we found a gradient of behavioral specificities, including neurons that were only active during one behavioral region, neurons that were similarly active throughout the whole behavior space, and neurons that fired throughout multiple but not all behavioral regions (Fig. 2a,b).

**Figure 2.**
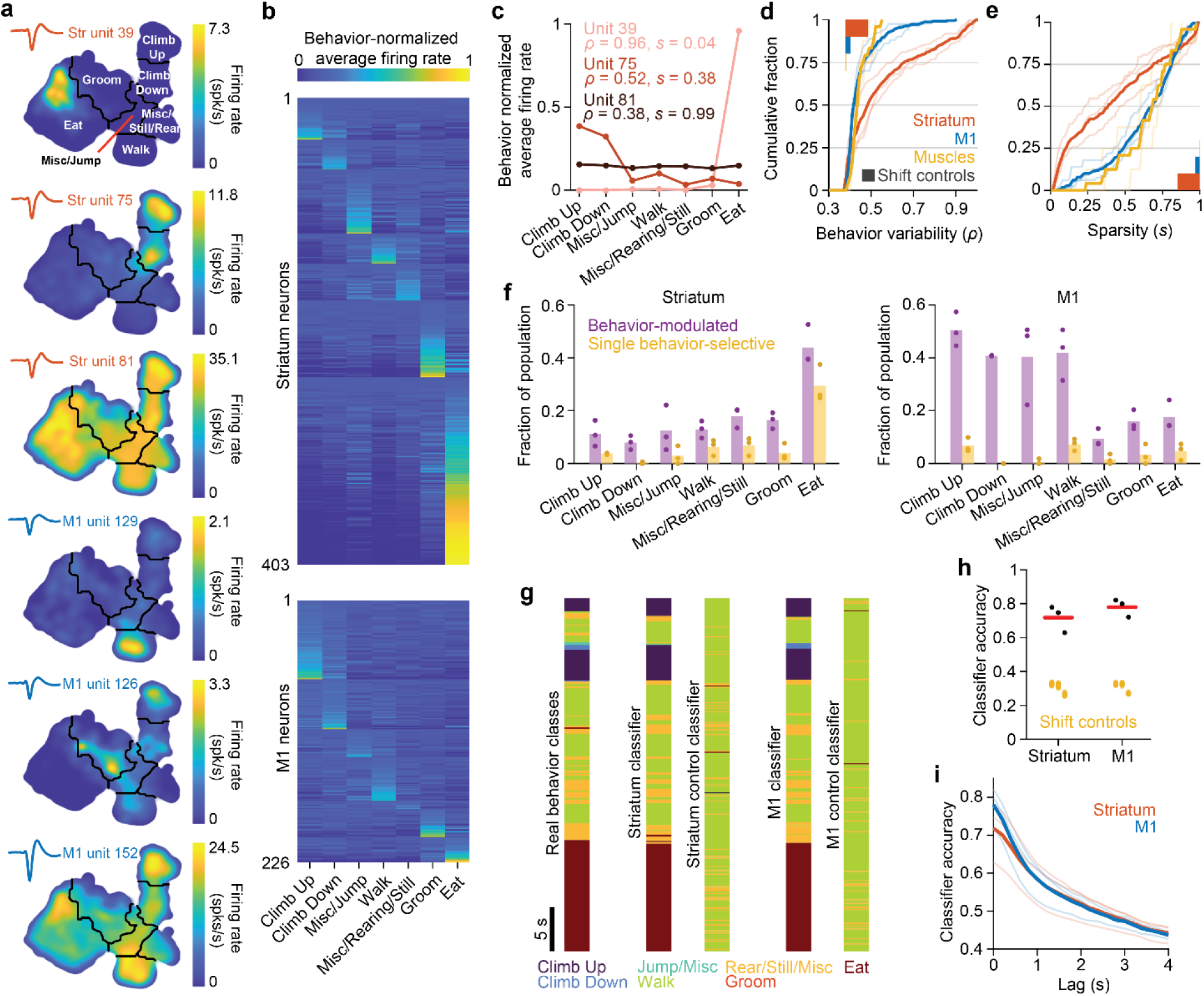
**| Behavioral specificity is broadly distributed in both areas, but stronger in striatum**. **a,** Activity from six example neurons overlayed in UMAP behavior space. **b,** Behavior-normalized average firing rates within each of the seven behavioral regions for all recorded neurons, sorted by the behavioral region with the highest activity. **c,** Behavior-normalized average firing rates across the seven behavioral regions from the three example striatum neurons shown in **a**. *ρ* is the value of the behavior variability metric and *s* is the sparsity index for each neuron. **d,** Cumulative histograms of *ρ* across the population for M1, striatum, and muscles. Bars on top indicate the 97.5^th^ percentile of the control distribution, calculated from circularly shifting the behavioral region labels relative to the neural or muscle activity. The striatal population exhibited larger *ρ* compared to the M1 population, 2 sample k-s test p < 1x10^-3^. For both **d** and **e**, thin lines represent individual mice, while the thick lines represent all mice combined. **e,** Cumulative histograms of *s* for both brain areas and the muscles. Striatum exhibits smaller *s* compared to M1, 2 sample k-s test p < 1x10^-3^. **f,** Fraction of neurons which were modulated by each of the behaviors (note that neurons can be modulated by multiple behaviors) and the fraction of neurons that were exclusively modulated by a single behavior. Dots denote individual mice, bars denote averages across mice. **g,** Behavioral region labels for an example time segment (left) along with the predicted labels from the striatum classifier (middle) or M1 classifier (right), and example control classifiers. **h,** Cross-validated classification accuracy for both classifiers. Each black dot represents a separate mouse, yellow dots represent the classifier accuracy using time shifted inputs for each animal. Difference was not significant between M1 and striatum, two-sided paired t-test p = 0.054. **i**, Classification accuracy versus time between neural activity and future behavior. Thin lines indicate individual mice, solid lines indicate average across mice.

To quantify the degree of behavior specificity in individual neurons, we defined a metric, the behavior variability (*ρ)*. To compute this, we first divided the mean firing rate of each neuron in each behavioral region by the sum of these mean firing rates across all regions, producing “behavior-normalized” mean firing rates. These normalized mean firing rates thus sum to one across all regions, facilitating comparison of how uniform or skewed the distributions are for different neurons. We then defined *ρ* as the L2 vector norm of these behavior-normalized rates. Neurons only active in one behavioral region would have a *ρ* of 1, while neurons with nearly the same firing rate for all seven regions would have a *ρ* near the minimum possible, 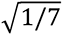 (Fig. 2c). We also employed another metric that quantifies behavior specificity independent of how the behavior space is parcellated into regions. We adapted the sparsity index (*s*) from analyses of place-dependent firing in the hippocampus^48,49^, replacing physical space with the behavioral space defined by UMAP. A lower sparsity value indicates that the activity of a neuron is confined to a smaller fraction of the total space (Fig. 2c).

Both metrics produced a broad range of behavioral specificities across neurons, finding both highly behavior-specific and highly behavior-invariant neurons (Fig. 2d,e). This was true for both M1 and striatum, although striatum exhibited greater overall specificity (average *ρ* = 0.449 for M1, 0.557 for striatum, average *s* = 0.608 for M1, 0.407 for striatum). Collectively, these results show that while we could uncover highly specific and highly invariant neurons, most neurons in both M1 and striatum lie along a continuum of behavioral specificity. Therefore, these motor areas do not adhere exclusively to a behavior-specific or behavior-invariant organization, although there are differences in the overall degree of behavior specificity between the two areas.

The greater overall specificity of striatal neurons was also apparent in the fractions of M1 and striatal neurons that were strongly modulated for individual behavior regions. We defined a neuron’s activity as modulated within a region if the activity was at least three standard deviations above the mean of a control distribution for the neuron’s activity in that region (derived through time-shifting behavior labels, see Methods). We found that both M1 and striatal populations contained neurons that were modulated for each behavior (Fig. 2f). On average, individual M1 neurons were modulated for more behaviors than were neurons in striatum (2.17 for M1, 1.23 for striatum, paired t-test *p* = 0.013). We then further defined a neuron as “single behavior-selective” if it was modulated for only one behavioral region. A higher percentage of neurons in striatum were single behavior-selective (53.4%) compared to M1 (24.2%, paired t-test *p* = 0.032, Fig. 2f).

We also investigated whether differences exist among different neuron types in each brain area. We divided neurons into wide-waveform (putative pyramidal neurons in M1, spiny projection neurons in striatum) and narrow-waveform (putative interneurons) populations (Extended Data Fig. 5a,b). Interestingly, narrow-waveform neurons in the striatum exhibited much lower behavioral specificity than wide-waveform neurons (Extended Data Fig. 5d,e), suggesting that they may play a more general role in motor control.

We next examined whether the neurons exhibiting high behavior-specificity were anatomically clustered. We observed a clear topography of eating-specific neurons in the striatum. Eating-related activity was localized to a ventral region, where many neurons suddenly activated at the onset of eating, while much of the rest of the striatum and M1 decreased in activity (Extended Data Fig 6a-c). This eating-specific activity was not reward related, as the food is placed into the arena irrespective of completion of any tasks. Nor was it pleasure-related modulation, as the majority of these neurons (mean = 84%) were not similarly active when the mouse received water during licking (Extended Data Fig 6d-f).

Finally, to test whether sufficient information was present in either neural population to distinguish between behavioral regions, we trained a classifier to predict the behavioral region given only the neural activity from M1 or striatum in each given mouse. Using a random forest classifier, we were able to accurately predict the correct behavioral region using either M1 or striatal activity as input (average accuracy was 0.78 for M1, 0.72 for striatum; Fig. 2g,h). Interestingly, classifiers were able to predict behavioral region even over 1 s into the future (Fig. 2i). Thus, while the striatum may exhibit greater behavior specificity at the single neuron level, M1 activity patterns are still distinguishable between behaviors at the population level.

### Neural activity is hierarchically organized across behaviors

Our findings above do not agree with an exclusively behavior-specific or behavior-invariant organization in M1 and striatum, as behavior specificity varied widely across neurons. We next sought to understand how activity patterns across M1 and striatal populations compared between behaviors. First, we examined whether neurons that were active during one behavior also tended to be active during a different behavior (Fig. 3a). If two behaviors share similarly active neurons, we would expect to see a clear correlation in average firing rates during those behaviors. Across all behavior pairs, we observed a range of correlation values, with certain behavior pairs, like climbing up and climbing down, exhibiting high similarity, while other behavior pairs, like climbing up and eating, exhibited much lower similarity (Fig. 3b).

**Figure 3.**
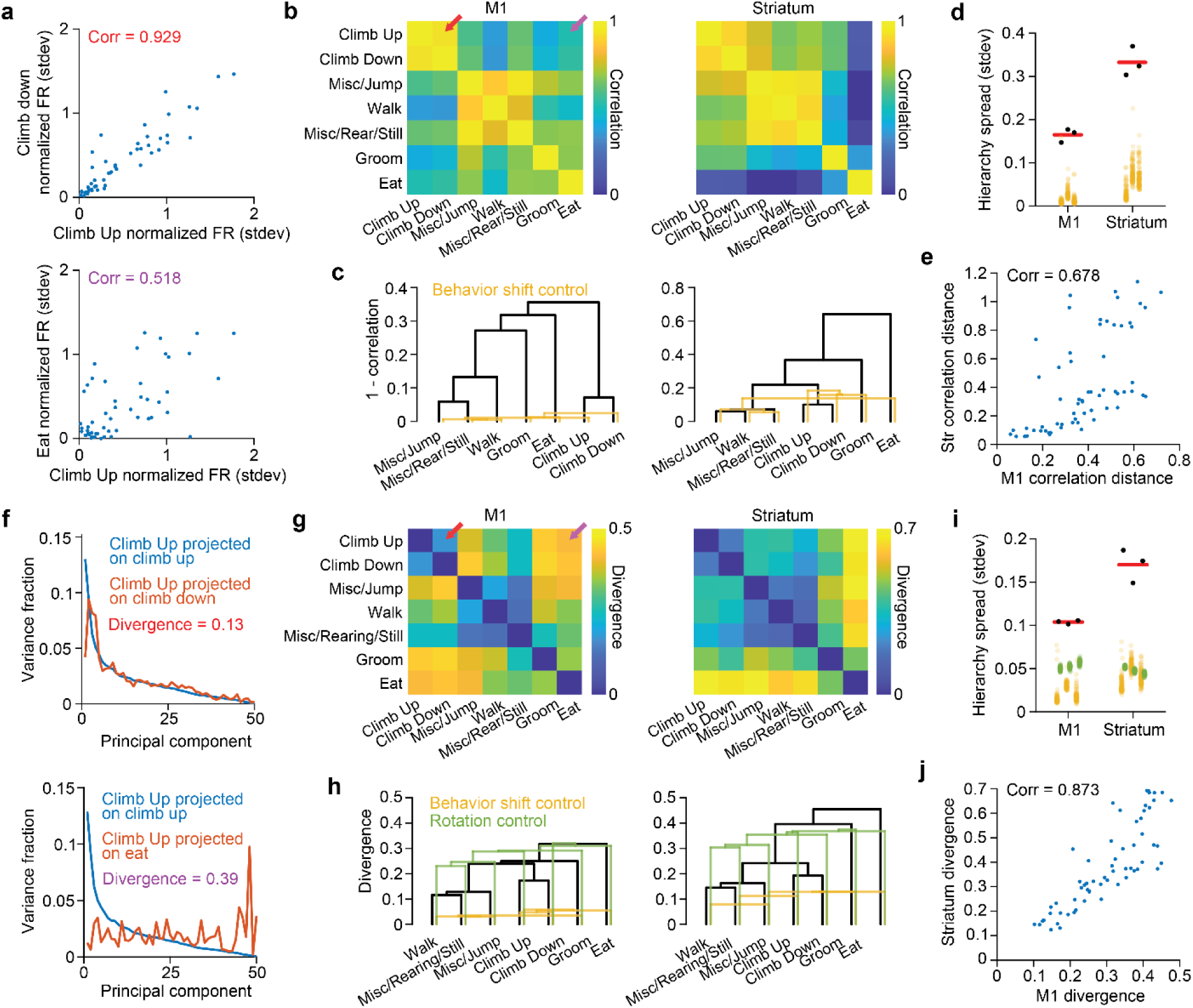
| Neural dynamics are hierarchically organized across behaviors. a, Example plots showing average firing rate, normalized by standard deviation, during climbing up against climbing down (top) or against eating (bottom) for all neurons in one mouse. The value in the inset indicates the Spearman correlation computed using all neurons. **b,** Spearman correlations for all pairs of behaviors in one mouse. Colored arrows indicate the corresponding behavior pairs shown in **a**. **c,** Dendrograms where the distance between behaviors was defined to be 1 - correlation. Yellow indicates one example dendrogram after circularly shifting the behavior labels. For both **c** and **h**, the example was the shift corresponding to the median of the 100 control hierarchy spreads. **d,** Hierarchy spreads for M1 and striatum. For both **d** and **i**, black dots indicates individual mice, red bar indicate average across mice. Yellow dots correspond to the 100 behavior shift-controls for each mouse. **e,** The correlation distance in M1 versus the correlation distance in striatum for all behavior pairs combined across all mice. Value in the inset indicates the Spearman correlation. **f,** Examples showing the variance captured across each principal component when the same neural activity during climbing up is projected into different PCA spaces computed during different behaviors. **g,** Divergence for all pairs of behaviors. Colored arrows indicate the corresponding behavior pairs shown in **f**. **h,** Dendrograms for the divergence across behaviors. Yellow is one example control dendrogram after circularly shifting the behavioral labels, green is one example control dendrogram after random rotations of the PCs. **i,** Same as **d**, except for dendrograms defined using divergence. Green dots denote 100 random rotation controls. **j,** Same as in **e** except for divergence rather than correlation distance.

To characterize the structure of this activity similarity across all behavior pairs, we employed hierarchical clustering, which groups elements together based on similarity defined by a distance metric^41^. Hierarchical clustering is commonly used in the analysis of gene expression^51–53^ or evolutionary relationships^54^. Here we used one minus the correlation value as the distance and illustrate the results using dendrograms (Fig. 3c). Clustering in this way produced relatively deep hierarchies: highly correlated behavior pairs are grouped together at lower levels, and behaviors exhibiting progressively lower correlations are connected at increasingly high levels. To determine whether the depth of these hierarchies arose simply due to chance, we generated control dendrograms by shifting the behavior labels in time. We measured the depth for both the observed and control dendrograms using the standard deviation of the correlation distance across all behavior pairs (Fig. 3d). For both brain regions and all animals, the hierarchy spread was significantly above chance (p = 0.0099 for all animals). Thus, the activity similarity between behavior pairs exhibited a hierarchical organization.

Given either a behavior-specific or behavior-invariant organization, we would not expect to see the emergence of a deep hierarchy, as the behaviors would have either all high or all low correlation distances, respectively. Indeed, the behavior label-shifted controls, which mimic behavior invariance, had similar neural activity and low distances across all behavior pairs, yielding a low spread and a lack of appreciable hierarchy (Fig. 3c,d yellow). Interestingly, we also found that hierarchies were similarly organized in M1 and striatum, as the correlations for behavior pairs were themselves correlated between the two regions (Fig. 3e).

However, average firing rates do not capture the interaction dynamics between neurons within a population^55,56^. We therefore sought to examine the degree of similarity across behavioral regions in terms of neural activity covariation. To characterize this covariation within each behavioral region, we applied principal component analysis (PCA) to the matrix of neural firing rates for time points falling within each region (neurons x time). This resulted in seven sets of PCs that reflect the activity covariation within each behavior region. To measure the similarity of the covariation between behaviors, we defined a divergence metric (Fig. 3f). This metric quantifies how similarly the activity variance is distributed across each behavior region’s PCs when the activity from one region is projected onto the PCs from two different behaviors (see Methods). Behaviors with similar covariation would have similar distributions of variance across PCs, while dissimilar covariation would lead to very different distributions of variance across PCs. This approach extends the subspace alignment index used previously to quantify the degree of overlap between low-dimensional subspaces in earlier studies^13,57^. Our divergence metric uses all PCs, not a subset.

Applying hierarchical clustering to divergence values for all behavior pairs, we found that activity covariation is also hierarchically organized across behaviors in both M1 and striatum (Fig. 3g-i). Some behavior pairs had low divergence, while others had high divergence. The spread of divergence-based dendrograms was significantly different from a null hypothesis of behavior invariance, as shifting the behavior labels in time resulted in similar covariation for all behavior pairs and low spread (Fig. 3h,i yellow, Extended Data Fig. 7a). The spread in divergence values was also significantly different from a null hypothesis of behavior specificity, as randomly rotating all PCs before computing the divergence resulted in high divergence for all pairs, and low spread (Fig. 3h,i green, Extended Data Fig. 7b). The hierarchies were again similarly organized in M1 and striatum (Fig. 3j). We obtained similar results using both the subspace alignment index of previous studies^13,57^, and the first principal angle between subspaces^28,58^ rather than divergence (Extended Data Fig. 7c-h). We also found similar hierarchical organization when using only time points from the human-labeled behavior classes rather than the seven behavioral regions defined via watershed segmentation (Extended Data Fig. 8).

To assess whether these hierarchies match those of the muscle activity itself, we computed the divergence between behavior pairs using muscle activity instead of neural activity. Unsurprisingly, similarity in muscle activity is also hierarchically structured, with some behavior pairs exhibiting low divergence and others exhibiting greater divergence (Extended Data Fig. 9a,b). However, the muscle activity divergence was only weakly correlated with cortical and striatal activity divergences (Extended Data Fig. 9c,d), especially compared to the correlation between cortex and striatum (Fig. 3j). Thus, pairs of behaviors that exhibit high divergence in muscle activity would not necessarily exhibit high divergence in cortical or striatal activity. Therefore, the neural activity hierarchies we observe are not expected simply because of differences in the muscle activity patterns across behaviors.

Taken together, these results further support our finding that neural activity in neither brain area is strictly behavior-specific or behavior-invariant. Rather, it is hierarchically organized across behaviors, with similar organization in M1 and striatum. We note that the term “hierarchy” we describe here is based on varying degrees of similarity between neural activity patterns^30,33,59^, and so is distinct from previous usages in describing feedforward organization among motor system regions^60,61^ or motor circuits subserving distinct functions^62,63^. While it is sometimes straightforward to assign categorical labels to different dendrogram branches after hierarchical clustering, what distinguishes the branches in our neural activity hierarchies is not always immediately apparent. Some behaviors, such as climbing up and climbing down can be thought of as subdivisions of a broader category, but other branches may be less intuitive to classify.

### Muscle activity is robustly encoded in M1 but correlations with neural activity vary across behaviors

Although neural activity does not appear to be behavior-specific or behavior-invariant in general, the activity components that correlate closely with muscle activity could themselves still be behavior-specific or behavior-invariant. This could result if muscle-correlated activity only constitutes a small fraction of the overall activity. We therefore sought to examine how muscle- correlated neural activity varies throughout behavior space.

First, we assessed the degree to which activity in M1 and striatum encodes muscle activity. We examined the time periods around muscle activity onsets, defined as periods where muscle activity was near baseline for at least 150 ms before increasing above a threshold (Fig. 4a). The average population firing rates around these onsets changed robustly in M1, but minimally in striatum (average increase over baseline was 0.830 spk/s for M1, 0.014 spk/s for striatum, Fig. 4b).

**Figure 4.**
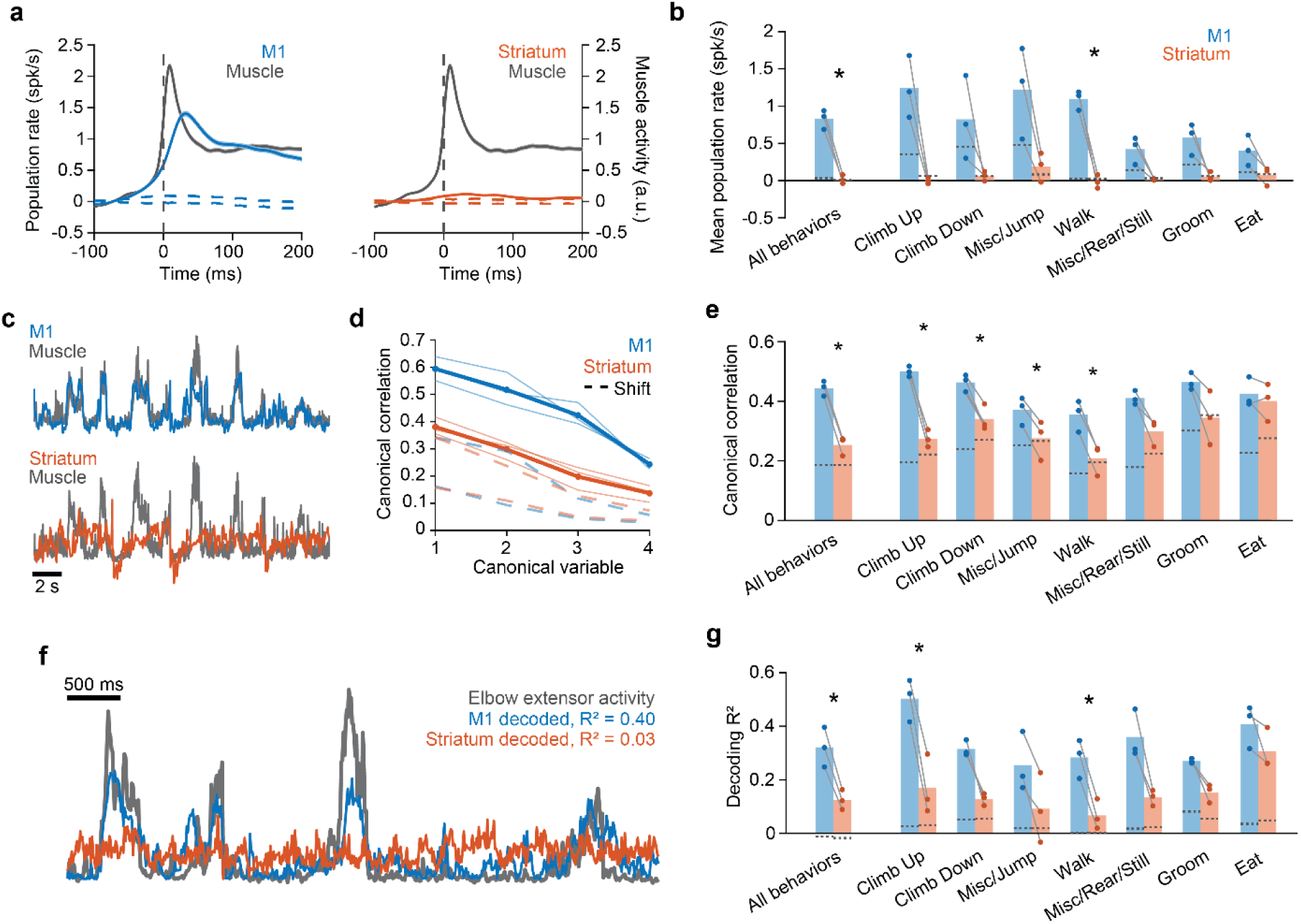
| Muscle activity correlations. a, Average population activity for cortical (left) or striatal (right) neurons triggered on muscle activations across all behaviors in one mouse. Activity was normalized to pre-trigger baseline. Dotted lines indicate the 2.5% and 97.5% percentiles from time shifting neural activity relative to muscle activity. **b,** Average population activity during the 200 ms after muscle activation triggers for all behaviors combined or each behavior individually. For **b**, **e**, and **g**, dots represent individual mice, bars represent the average across all mice, and dotted lines indicate the 97.5^th^ percentile of shift controls. Stars indicate statistical significance (two-tailed paired t-test, with Benjamini Hochberg multiple comparisons correction, FDR < 0.05). **c,** Example time series during climbing up of the first canonical variable for M1 (top) and striatum (bottom) along with the corresponding muscle canonical variable. **d,** Canonical correlations between neural activity and contralateral forelimb muscle activity during all behaviors combined for individual mice (thin), or average across mice (thick). Dotted lines indicate the 2.5% and 97.5% percentiles from time shift controls. **e,** Average canonical correlation across all four canonical variables. **f,** Example time series of right elbow extensor activity during climbing up, along with activity predicted from decoders using M1 or striatal inputs. **g,** Decoding performance (weighted average across all contralateral forelimb muscles).

Next, we performed both canonical correlation analysis (CCA), and decoding analysis using either M1 or striatal activity. Consistent with the muscle-triggered results, CCA revealed that M1 activity was more strongly correlated with muscle activity compared to activity in striatum, both combined across all behavior regions and for each individual region (Fig. 4c-e). Additionally, Wiener cascade decoders could more accurately reconstruct the time course of muscle activity from M1 activity compared to striatal activity (mean R^2^ = 0.32 for M1, 0.13 for striatum, Fig. 4f,g). Thus, the strengths of muscle activity representation in M1 and striatum are markedly different.

We then examined how the correlation between neural and muscle activity varied across behaviors. Because the striatal population encoded muscle activity only very weakly, we limited our analysis to just M1 neurons. For each of the seven behavioral regions, we computed the Pearson correlation between each neuron’s firing rate and the activity of the four contralateral forelimb muscles (Fig. 5a). To determine whether neurons that were correlated to muscles during one behavior remained similarly correlated during another behavior, we calculated the Spearman correlation across all neuron-muscle pairs, for each pair of behaviors. Across all behavior pairs, we observed a range of stability in the neuron-muscle correlations (Fig. 5b). Certain behavior pairs like climbing up and climbing down maintained stable correlations, while other behavior pairs like climbing up and eating exhibited more variable correlations (Fig. 5a). As for the neural activity in general, the patterns of neural activity related to muscles were hierarchically organized, having a hierarchy depth much greater than controls (Fig. 5b-c).

**Figure 5.**
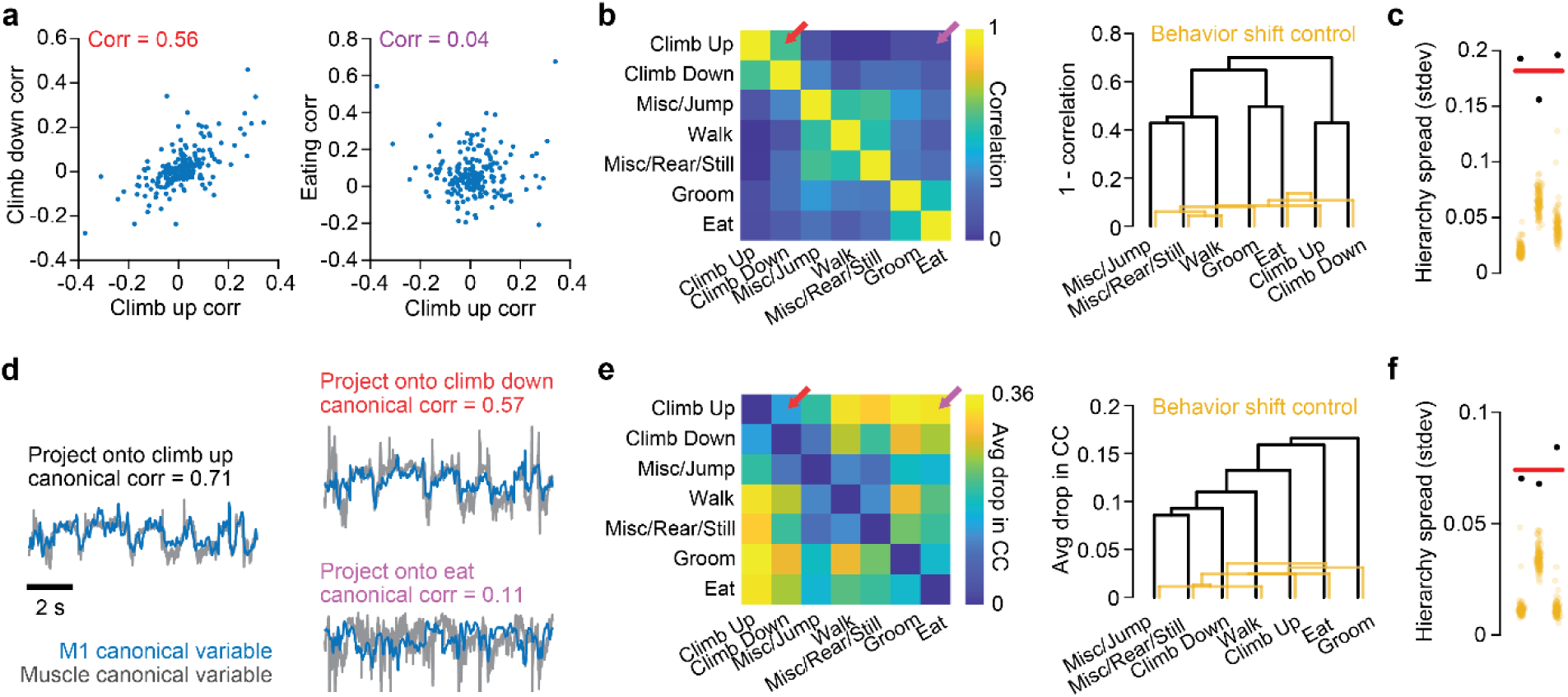
**| Neural-muscle correlation stability across behaviors**. **a,** Correlations for all neuron-muscle pairs during climbing up against climbing down (left) and during climbing up against eating (right) in one example mouse. The value in the inset indicates the Spearman correlation computed using all neuron-muscle correlations. **b,** Spearman correlation for all pairs of behaviors (left), and the dendrogram representing the hierarchy of the correlation distance, in one mouse. Colored arrows indicate the corresponding behavior pairs shown in **a**. For both **b** and **e**, yellow indicates an example control dendrogram after randomly time shifting behavior labels. **c,** Correlation-based hierarchy spreads for each mouse (black dots) and the average across mice (red bar). For both **c** and **f**, yellow dots indicate the hierarchy spread for the shift controls in each mouse. **d,** Time series of the top neural and muscle canonical variable during climbing up (left), and the same neural and muscle activity projected using the CCA weights computed during climbing down (right, top) or during eating (right, bottom) for one mouse. The values indicate the correlation between the top canonical variables when projected this way. **e,** Drop in correlation, averaged across all four canonical variables, for all pairs of behaviors (left) and the dendrogram of the drop in correlations across behaviors (right) in one mouse. Colored arrows indicate the corresponding behavior pairs shown in **d**. **f,** CCA-based hierarchy spreads for each mouse (black dots) and the average across mice (red bar).

We also examined the variability of muscle correlations on a population level using CCA. We computed canonical vectors within each behavior region, then calculated the average drop in canonical correlation when data from one behavior is projected onto the canonical vectors from another behavior (Fig. 5d). We again observed a range of similarities that formed a hierarchy across behaviors (Fig. 5e,f). Taken together, these analyses indicate that the patterns of correlation between M1 and muscle activity are not fixed or purely behavior-specific, but show varying similarity across behaviors.

### Striatal inputs are sufficient to drive modulation of cortical dynamics

In one prominent model of action selection, the striatum selectively inhibits or disinhibits downstream motor circuits to promote execution of particular behaviors^35,36,64^. The strong behavior specificity of individual striatal neurons but weak relation to muscle activity appears to conform to this model. However, in M1, rather than distinct neuronal subpopulations corresponding to different behaviors, we found graded shifts in covariation patterns. To update this model in light of our observations, we hypothesize that striatum is able to drive graded modulation of population dynamics in descending control areas like M1.

To test this, we utilized a recurrent neural network (RNN) which has been shown to faithfully capture the activity dynamics of motor cortex during movement^65^ (Fig. 6a). Because RNNs are generally less accurate at reconstructing long time series, we restricted our analysis to 300 ms windows surrounding specific movement events. For six different behaviors, we identified instances of characteristic limb movements, such as the hand strike during locomotion, or when the mouse initiated a reach during climbing. We then computed the average activity of each contralateral forelimb muscle across all instances of these movements (Fig. 6a,b, gray) and trained an RNN to reconstruct the average muscle activity given the observed average activity of striatal neurons as inputs (Fig. 6b, middle left). As a positive control, we used a unique constant input for each behavior (Fig. 6b, top left); as a negative control, we used averages of time-shifted striatal neuron activity as inputs (Fig. 6b, bottom left).

**Figure 6.**
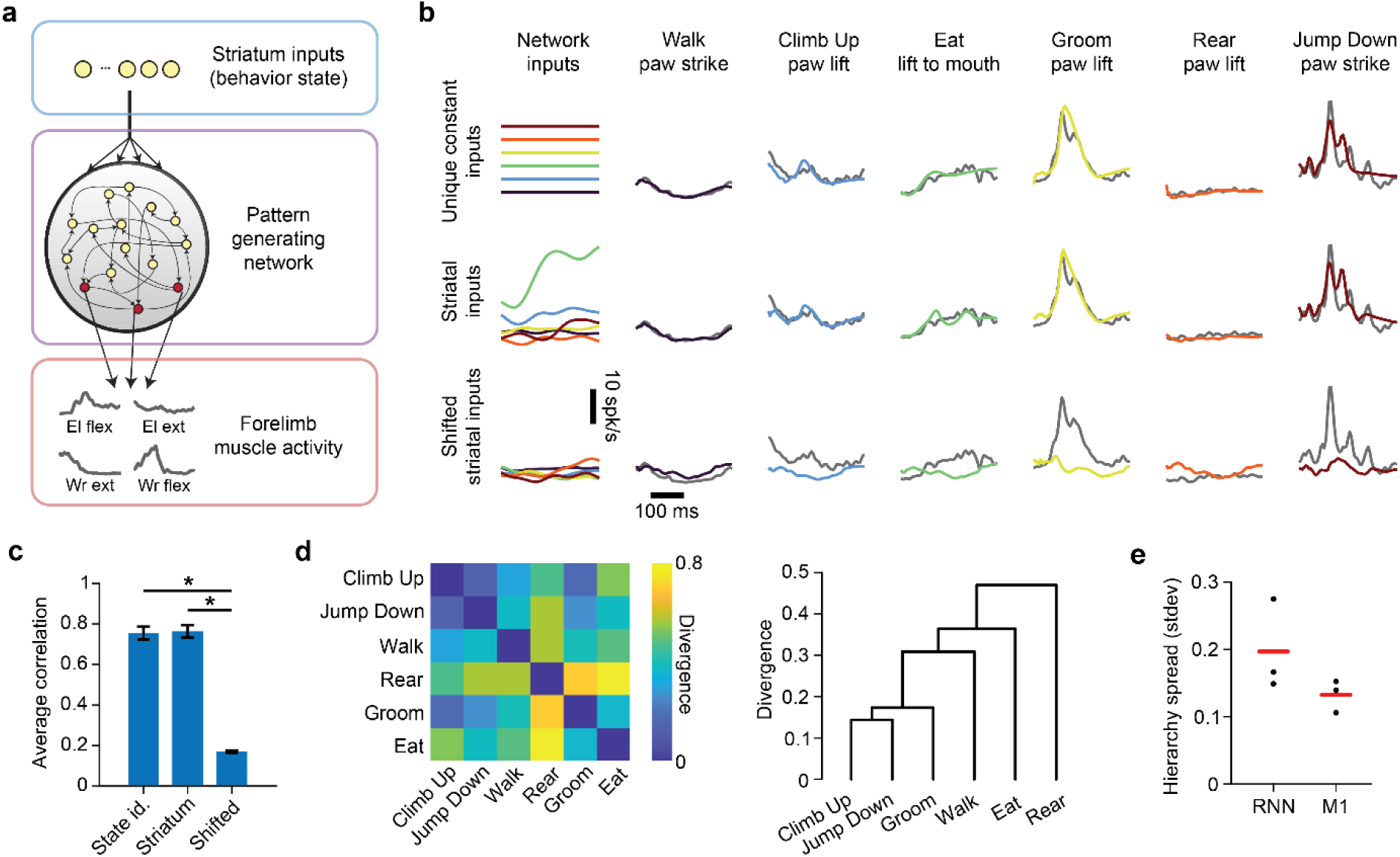
| Network modeling. a, Illustration of the RNN used to generate muscle activity. **b,** Example model inputs and outputs. Left-most column illustrates the time-series inputs to the network (for the striatal and shifted striatal inputs, the plots show the activity of one example neuron). Remaining columns show example activity from one muscle (gray) and the outputs of the RNN (colored) for each of the different behaviors. **c,** Average correlation between the actual muscle activity and the model outputs across all behaviors and all muscles of all mice. Error bars denote SEM, Wilcoxon rank sum test, p < 10^-3^ for comparisons to shifted controls, p = 0.868 for comparing state identity and striatum inputs. **d,** Example subspace divergence across pairs of behaviors in the activity of the units from the RNN model for one mouse. **e,** Divergence-based hierarchy spread for both the RNN units and the recorded units from M1. Two-tailed paired t-test p = 0.188.

We found that observed striatal activity is able to drive the network to accurately produce the observed muscle activity, with accuracy similar to that of the constant inputs (mean Pearson correlation of 0.76 for striatal inputs and 0.75 for constant inputs), while the shifted striatal inputs could not (average correlation of 0.17, Fig. 6b,c). We then examined the activity of the RNN units and found that it displayed a range of divergence values across behavior pairs, forming a hierarchy with similar spread as the hierarchy of actual M1 activity (average spread of 0.20 for RNN units and 0.13 for cortical units, Fig. 6d,e). Thus, the activity in the striatum is sufficient to drive a network to modulate through a range of activity dynamics to produce varied patterns of muscle activity across behaviors.

## Discussion

In this study, we introduce a novel paradigm that enables examination of motor system activity across a broad range of naturalistic behaviors. We demonstrate that activity in forelimb M1 and striatum does not exclusively exhibit a behavior-specific or behavior-invariant organization, but spans a continuum of specificity that is organized hierarchically across behaviors. This hierarchy is present in the activity of individual neurons, population-level activity covariance, and muscle- correlated activity, and is similarly organized in M1 and striatum. Despite this commonality between areas, we found marked differences in the properties of M1 and striatal activity, with only M1 neurons strongly encoding muscle activity but striatal neurons exhibiting greater behavior specificity. Finally, our network modeling revealed that specificity in striatal activity is sufficient to modulate the dynamics of downstream muscle-controller circuits, such as those in M1, creating hierarchical neural dynamics that can produce muscle activation patterns for a range of behaviors.

Neural dynamics hierarchy constitutes a framework for understanding the variation of neural dynamics across tasks. Hierarchy here refers to an organization defined by hierarchical clustering, as has previously been used in analysis of gene expression^51,52^, evolutionary relationships^54^, and motor behavior types^30,33,59^. Here the hierarchy is based on the relative similarity of neural dynamics between behaviors, where behavior pairs exhibit progressively dissimilar dynamics the further up in the hierarchy they are linked. This hierarchy can account for the seemingly contrasting reports of either behavior-specific neurons and dynamics^6,7,9–12^, or similar dynamics across different behaviors^26,28,66^. Previous studies have focused on only a small number of behavior types^10,26,28,30^, which in our framework could lead neural activity to appear either very similar or very different depending on where within the hierarchy the behaviors lie. Individual neurons that appear behavior-specific in some paradigms may lie near one extreme of the specificity distribution, or may activate during other, unobserved behaviors. Only by examining neuronal populations across a broad enough range of behaviors can the hierarchy be well resolved.

Movements themselves are naturally hierarchical^32,43,59,67,68^ – different movements can involve kinematics or muscle patterns that exhibit differing degrees of similarity. Indeed, we were able to derive a hierarchy based on muscle activity. However, the similarity based on muscle activity divergence was only weakly correlated with similarity based on cortical or striatal activity divergence. This suggests that the neural activity hierarchy we observed is not a simple consequence of changes in muscle activity patterns. Furthermore, we found that neurons did not maintain the same correlation with muscles across all behaviors. Consequently, a hierarchical organization of muscle activity does not imply that the same organization will emerge in the corresponding neural activity. It is therefore unlikely that the hierarchies we observe arise simply from the similarities and differences in the movements themselves.

This neural dynamics hierarchy may reflect the various combinations of computational elements associated with different behaviors, known as compositionality^69,70^. For example, reaching for a handhold during climbing or the grid floor during walking may both involve a sensorimotor transform of an external target location, but such a computation is not necessary during grooming. Different behaviors may involve varying degrees of preparation^71^, attention^72,73^, or sensory feedback processing^74,75^. These computational elements may each rely on a distinct neural activity subspace^13,57,71,76–78^; the varying combination of elements during each behavior would then give rise to the varying similarity in neural dynamics we observe. Compositionality may reflect a tradeoff between the fitness benefits of reusing computational elements (generalization) or employing task-specific elements (specialization)^29^.

One striking feature of our data was the relatively strong behavior specificity in striatum. In particular, we found distinct eating-specific ensembles, consistent with previous reports of functional groups in striatum^31,79–81^. In contrast to the substantial overlap of corticostriatal projections from different cortical regions, at least in data aggregated across mice^82^, these eating-specific ensembles appear to be localized to a discrete region within the ventral striatum. This could reflect a behavior-based topography within the striatum^83^ with relatively discrete boundaries between regions. Given the association of ventral striatum with reward-based learning^84,85^, one might believe that the responses in these neurons are related to reward^86–88^ rather than movement^89,90^. However, the timing of food placement was random and not tied to any task completion, and the majority of these neurons were not similarly active during licking, which should be more rewarding since the animals were water restricted.

The much greater behavior selectivity of striatal activity compared to M1, and its nearly absent correlation with muscle activity at high temporal resolution seem at odds with the strong anatomical and functional connectivity between M1 and striatum^82,91–93^ – motor and somatosensory cortices are the largest source of input to striatal output neurons. Even corticospinal neurons, which encode motor commands^94,95^, send abundant collaterals to striatum^93^. It remains to be seen how striatal activity avoids muscle correlation, diverging sharply from its primary input. These distinctions between M1 and striatum activity also contrast with recent evidence for the encoding of movement kinematics in striatal activity^96–99^. These recent findings have sparked debate over whether striatum implements high-level functions like behavior selection, as in traditional models, or lower-level control of on-going movement execution^39,100,101^. Our direct comparison of striatal and cortical activity across a broad range of behaviors revealed differences that align with a role in high-level function. However, because our recordings were limited to just a single narrow track through striatum; it remains possible that execution-related circuits may lie in a spatially distinct location^96,102^.

Our results could point to a more nuanced interaction between action selection and effector- control circuits, which has traditionally been modeled as involving gross activation and suppression of muscle pattern-generating circuits related to different movements^35–39^. In M1, we observed short-latency influence on muscles and encoding of muscle activity, but cortical neurons did not express strong single-neuron selectivity for behaviors. Rather, we found that the coordinated activity across the population shifted to varying degrees as the mice engaged in different behaviors. We therefore propose that rather than implementing action selection by selectively activating different subpopulations corresponding to different behaviors, the striatal influence on M1, through motor thalamus^103–106^, could drive cortex to adjust its population-level activity dynamics as animals perform different actions^74,107,108^. The implied role of thalamocortical projections here aligns with emerging views of their involvement in cognitive control^109,110^. Our modeling results support such a role for striatal influence, although we note that these results do not preclude the possibility of additional inputs driving the modulation of population dynamics in M1. Indeed, it is likely that a combination of the highly behavior-specific inputs from the basal ganglia, and inputs related to the unique demands of individual behaviors, are what produces the changes in activity dynamics we observe in M1.

## Methods

### Experimental animals

A total of 41 adult male mice were used including those in early experimental stages to develop the new paradigm and establish methodology. Strain details and number of animals in each group are as follows: 14 VGAT-ChR2-EYFP line 8 mice (B6.Cg-Tg(Slc32a1-COP4*H134R/EYFP) 8Gfng/J; Jackson Laboratories stock #014548) and 27 C57BL/6J mice (Jackson Laboratories stock #000664). All mice used in experiments were individually housed under a 12-h light/dark cycle in a temperature- and humidity-controlled room with *ad libitum* access to food and water, except during experiments. At the time of the measurement reported, animals were 13–22 weeks old. Animals weighed approximately 22–28 g. All animals were being used in scientific experiments for the first time. This includes no previous exposures to pharmacological substances or altered diets. All surgical and experimental procedures were approved by the Institutional Animal Care and Use Committee at Northwestern University.

### Arena paradigm

The arena was housed in an illuminated room and contained an acrylic base with four slots cut out to allow insertion and removal of different walls and different floors (Fig. 1a). Walls consisted of 12” high x 18” wide x 3/32” thick acrylic pieces (McMaster-Carr) containing climbing handholds and laser cut slots to serve as insertion sites for water ports. Handholds were made from copper mesh cut into 8 mm diameter circles and attached to the wall with clear epoxy. Handhold locations were randomly spaced, with horizontal and vertical spacings ranging between 0.35 to 1.75 inches. We did not place any handholds within the top 2” of the walls; however, some mice were still able to climb out of the arena. For these mice, we would additionally place an acrylic lip above the walls, overhanging the top inner edge by 1.5”.

The floor was 18” long x 18” wide x 1/4” thick. It contained a large grid region where most of the floor was removed, leaving only thin 3/32” wide handholds for walking. The spacing of this grid was also randomly generated, with a range between 0.5” and 1.5”. The floor also contained several solid flat regions (dimensions 1.5” x 1.5”, 3” x 3”, 2” x 8”, and 8” x 8”) allowing for overground walking without requirements for accuracy. Finally, we added an aversive 4” x 8” “spike pit” region which consisted of an array of 3D printed plastic cones with a small 1.5” x 1.5” flat platform in the middle. The floor was raised 0.4” above the arena base so that mice walked along the grid. During training and across different mice, the walls were intermittently swapped out with walls containing different handhold patterns, but throughout all experimental recording sessions within each mouse, the wall patterns were kept constant. The floor pattern was kept constant throughout all experiments of all mice.

To motivate the mice to explore the full extent of the arena, we placed up to four water ports at different locations. These ports consisted of hollow metal tubes attached to the end of flexible plastic tubing (C-Flex, Masterflex) which ran to solenoid valves (AT012IC, NResearch) housed above the arena in an electrically shielded enclosure. The valves are under electronic control to allow water to flow from a water reservoir to the port. The metal tubes were also connected through a wire to capacitive touch sensors (AT42QT1011, Sparkfun) which detected when the mouse made contact with the port. The port was attached to a 3D printed twist-and-lock design (Fig. 1a), which allowed quick placement into, or detachment from, various slots in the walls. A DAQ (PCI-e-6323, National Instruments) controlled by MATLAB running on a control computer (Extended Data Fig. 1c) detected touch events from the sensors and sent signal pulses to the solenoid valves to briefly open them.

Mice were allowed to behave freely within the arena. To encourage continuous exploration, only one water port was selected to be active at any time. Each time a lick was detected on the active water port, 3 µL of water was dispensed. Each port only dispensed 18 µL in total before deactivating, and one of the other ports would be randomly selected to become the new active port (Extended Data Fig. 1b). Each session lasted until mice received a total of 350 dispensations, up to a maximum of 120 minutes. A MATLAB program controlled the activation and selection of ports. To promote the expression of oromanual behavior, a sunflower seed was manually placed inside the arena every 5-10 minutes.

### Training

Within the first session of being introduced to the arena, mice naturally learned to walk across the grid floor and jump across the spike pit without any behavior shaping. They also naturally performed ethological behaviors such as rearing, eating, and grooming. However, most mice did not immediately use the wall handholds to climb. We therefore developed a behavior shaping procedure to encourage mice to climb for rewards over the course of two weeks.

Mice were first placed on a water schedule where they received 1 mL of water each day. After at least three days, they were placed inside the arena and introduced to the water ports by placing a single port at an easily accessible location near the ground. The port would dispense 18 µL of water before entering a time-out period of 2-5 seconds where they would not dispense water in response to licking. Mice quickly learned to associate the port with receiving water reward.

We next used a specially designed training wall consisting of climbing handholds that were closely spaced adjacent to each other and numerous port insertion slots spanning the height of the wall. The dense spacing of these handholds allowed mice to easily grab the next handhold without requiring a specific hand placement location. The height of the water port would be incrementally increased until mice were able to climb to the topmost position. After mastering this “easy” variant, mice then graduated to the “difficult” climbing pattern where the spacing of the handholds reflected the actual sparse spacing during recording sessions. Like before, the water port would start at the lowest position, then gradually be raised until mice were able to reach the highest position on the wall.

After the mice were able to consistently climb to the top, they would then be acclimated to the presence of multiple water ports. We first introduced a second water port that would always be placed close to the location of the first port. This port would then be gradually moved to further locations and to different walls. After mice learned to move between the two ports, they were introduced to a third, and finally fourth water port at various locations in the arena. Mice were considered proficient when they were able to acquire 350 dispensations across 4 different ports within a single session. During experimental sessions, some animals would occasionally have difficulty locating port locations when first entering the arena. In these cases, we would leave a trail of water drops from the port to the bottom of the wall using a syringe to help the animals find the port for the first time. Animals were maintained on the same water schedule and given *ad libitum* access to food throughout all experimental sessions.

### EMG recordings

EMG electrodes were fabricated in house following standard protocols established in previous studies^13,42,111^. Each electrode set contained 8 twisted pair wires made from 0.001″ braided stainless steel (793200, A-M Systems) knotted together. For wire pairs implanted into the forelimbs, the insulation was removed 1 to 1.5 mm from the knot on one of the wires and 2 to 2.5 mm away from the knot on the other wire, exposing the bare metal (electrode sites). For hindlimb wires, the insulation was removed 1 to 2 mm and 3 to 4 mm from the knot instead. The ends of the wires above the knot were soldered to a 16-pin miniature connector (CLP-112-02-F-D, Samtec). The amount of wire between the knot and the connector was 3.5 cm for the elbow flexors and extensors, 4.5 cm for the wrist flexors and extensors, 8 cm for knee extensors, and 9 cm for ankle flexors and extensors. The ends of the twisted section of the wires were inserted into a 27- gauge hypodermic needle that was then crimped to secure the wire inside. The needle enabled insertion into muscle.

All animals had electrodes implanted in the extensors and flexors of the elbow and wrist of both forelimbs as well as the ankle flexor of the contralateral hindlimb. One animal (mouse D020) had electrodes additionally implanted in the ankle extensor of the contralateral hindlimb while all other animals had electrodes implanted in the knee extensor of the contralateral hindlimb instead.

At the start of the insertion surgery, the head, neck, limbs and back of the mice were shaved. Mice were anaesthetized with 1-3% isoflurane and incisions were made in the skin above the muscle insertion sites and at the base of the skull. Wires were tunneled under the skin from the neck to the insertion site. Using the needle, the wires were then inserted into the muscle, pulling the metal contact sites into the belly of the muscle. The wire at the distal end of the muscle was knotted and the needle and excess wire was cut off. Finally, the incisions were sutured closed and the connector was attached to the base of the skull using dental cement (Metabond, Parkell). Depending on the experimental group, mice would then be implanted with either a ferrule guide or a Neuropixels probe in the same surgery (see Optogenetic inactivation and Neural recording sections below).

Before experimental sessions, mice would be briefly anesthetized with 3% isoflurane to allow attachment of an interconnect cable to the EMG connector on the skull. The other end of the interconnect cable was attached to a motorized commutator (CMTR-12/24-M-INTAN-NT, NeuroTek-IT) to allow mice to rotate without twisting the wires. Animals were allowed to recover from anesthesia for at least 30 minutes before being placed into the arena. The EMG signals were amplified, filtered (wideband, 1 Hz to 7.5 kHz) and digitized by a bipolar recording headstage (C3313, Intan Technologies) before being sent to a controller board (C3100, Intan Technologies) connected to the recording computer (Extended Data Fig. 1c). The EMG signal was acquired at 20 kHz using the Intan RHX Data Acquisition Software.

### Optogenetic inactivation

During surgery, after completion of the EMG electrode insertions, VGAT-ChR2-EYFP mice were moved to a stereotactic frame (Kopf Instruments) and the skull was fixed in place. A 2 mm diameter craniotomy centered at 0.25 mm rostral, 1.5 mm lateral relative to bregma was performed to expose the caudal forelimb area (CFA) in the left hemisphere. A silicone adhesive (Kwik-Sil, World Precision Instruments) was applied over the dura and a 3 mm diameter #1 thickness cover glass (64-0720, Warner Instruments) was placed on top. The edges of the coverslip were sealed to the skull using dental cement. A custom designed 3D printed ferrule guide was placed 1 mm above the brain over the craniotomy and secured to the skull using dental cement. The ferrule guide was positioned to allow delivery of a 2 mm diameter spot of light onto the exposed brain surface (Fig. 2a). Finally, the skin over the skull was secured to the dental cement with a tissue adhesive (Vetbond, 3M).

Before experimental sessions, a 240 µm diameter plastic core, 0.5 numerical aperture patch cable terminating in a 1.25 mm diameter zirconia ferrule (Doric Lenses) was secured to the ferrule guide using a set screw at the same time the EMG interconnect cable was attached. This patch cable ran to a fiberoptic rotary joint (FRJ_1x1_FC-FC, Doric Lenses) connected to the commutator. The other end of the joint was connected by a second patch cable to a 473 nm laser (PSU-III-LED, Opto Engine LLC). During inactivation experiments, a single 50 ms duration pulse of light at 10 mW/mm^2^ intensity was applied to CFA at random intervals between 1 to 1.3 seconds throughout the whole session as animals freely behaved in the arena.

### Neural recording

Mice were chronically implanted with Neuropixels 1.0 probes^47^ (Imec) following the methods modified from a previous publication^50^. First, the tip of the probe was sharpened using a microgrinder (EG-45, Narishige) to allow penetration through the dura. The probe was then attached to 3D printed head fixture pieces using cyanoacrylate and then coated in DiI stain (D282, Invitrogen) for histological visualization of the probe tract. We modified the fixture designs from the previous publication^50^ to account for the specific insertion angle and location in this study.

During surgery, after completion of the EMG electrode insertions, mice were moved to a stereotactic frame and a small <0.5 mm diameter craniotomy was performed above left CFA (centered at 0.5 mm rostral and 1.7 mm lateral relative to bregma). The probe was attached to a micromanipulator arm (uMp-4, Sensapex) and lowered 4 mm into the brain at a 10° angle in the coronal plane and at a rate of 4 µm/sec. The exposed brain was covered with silicone gel (3-4680, Dow Dowsil) and the fixture was secured to the skull with dental cement. A second craniotomy was made over the right visual cortex and a stainless steel ground wire was inserted into the brain. This wire was connected to the reference pad on the probe. The Neuropixels headstage was attached to the probe through a flex cable and inserted into a slot in the fixture. Finally, the skin over the skull was attached to the dental cement with tissue adhesive and the whole assembly was wrapped in Kapton tape (McMaster-Carr).

Before experimental sessions, the headstage was connected to the commutator through an interconnect cable. The other end of the commutator was connected to a PXIe acquisition card (Imec) in a PXIe chassis (NI PXIe-1071, National Instruments) which interfaced with the recording computer through a PXIe card (PXIe-PCIe8381, National Instruments, Extended Data Fig. 2c). Recordings were referenced to the probe tip, and data was acquired at 30 kHz using the SpikeGLX software (Janelia Research Campus).

### Video recording and behavior annotation

Video recordings were acquired at 41.55 fps from a side view of the arena with a 2.3 MP camera (BFLY-U3-23S6M-C, Teledyne FLIR). To synchronize the video, EMG, and Neuropixels acquisition systems, sync pulses were intermittently generated, which would flash an LED placed in view of the camera. These sync pulses were simultaneously sent to both the sync port of the Neuropixels acquisition card and an analog acquisition channel on the Intan recording board.

To identify time segments corresponding to different behaviors, we performed manual annotation of the video recordings. We defined 10 non-overlapping behavioral classes based on the following criteria: Climbing up – the mouse is on the climbing wall with all four limbs off the ground, and the body is only moving upwards.

Climbing down – the mouse is on the climbing wall with all four limbs off the ground, the body is moving downwards, and only downwards for the rest of the climbing bout.

Jumping down – the mouse dismounts from the wall by having all four limbs in the air (not contacting the wall or floor) for at least 1 frame. 3-5 frames before the mouse leaves the wall and after the animal hits the ground are included for each jumping down bout.

Jumping across – the mouse starts from and lands on the floor, and has all four limbs in the air (not contacting the floor) for at least 1 frame. 3-5 frames before the mouse leaves the ground and after the animal hits the ground are included for each jumping across bout.

Rearing – the mouse lifts both forelimbs off the ground while having both hindlimbs on the ground for at least 5 frames.

Still – the mouse does not move any of its limbs for at least 20 frames. Eating – the forelimbs of the mouse are in contact with the sunflower seed.

Grooming – the mouse is upright, and performs at least two cycles of cyclic movements with its forelimbs contacting either the face or the body.

Grid Walking – the mouse performs at least 2 continuous gait cycles with no pauses while all four limbs are only in contact with the grid sections of the floor.

Flat Walking – the mouse performs at least 2 continuous gait cycles with no pauses while all four limbs are only in contact with the flat sections of the floor.

Other miscellaneous behaviors not falling under these 10 classes were assigned to an “unlabeled” class. Additionally, there were frames where the mice could be subjectively associated with one of these behaviors but did not strictly fall under the defined criterion (for example the animal briefly performing one stroke on the face for grooming, or the animal briefly moves downwards during climbing before resuming ascent). These frames were also assigned to the “unlabeled” class.

### EMG data preprocessing

EMG signals were band-pass filtered between 100 and 1000 Hz, rectified, and enveloped by convolution with a 40 ms moving average box filter. We identified artifacts arising from contact with the capacitive lick ports by detecting when voltage levels crossed a threshold. We removed artifact timepoints and 100 ms before and 150 ms after artifacts from the data. Unless otherwise specified, we used EMG signals downsampled to 1 ms time bins. We divided the activity in each EMG channel by the standard deviation across the whole recording session for the analyses that used normalized muscle activity.

### Neural activity preprocessing

To remove spatially localized noise that was present across groups of electrode sites, we performed local median subtraction on each channel using the surrounding 10 channels above and below the channel; we found this was more effective than just performing a global median subtraction computed from all the channels. We also removed artifact time periods by removing data 100 ms before and 333 ms after time points where the voltage levels exceeded an artifact amplitude threshold (here 470 µV, based on inspection of artifact amplitudes) in the neural activity. We extracted putative single units from the Neuropixels recordings using automated spike sorting (Kilosort3^112^) followed by manual curation based on autocorrelograms and waveform PCA clustering. We excluded units which contained less than 100 spikes across the whole recording session.

We used both anatomical and electrophysiological properties to separate striatal and cortical populations. Based on our histology images, the thickness of cortex in CFA ranged between 1.2-1.4 mm. We observed a sharp drop off in the number of sorted units around this depth, corresponding to the white matter tract of the corpus callosum. For each animal, we computed the histogram of cell counts across depth using 100 µm bins (Extended Data Fig. 4c), and defined the M1-striatum boundary as the first depth bin after 1.2 mm with no identified units. Finally, we computed smoothed firing rates by summing the number of spikes in each time bin followed by convolution with a gaussian (10 ms standard deviation for 1 ms binned firing rates, 30 ms standard deviation for 10 ms binned firing rates).

To distinguish between putative interneurons and putative pyramidal or spiny projection neurons, we fit a two-class gaussian mixture model to the distribution of peak-to-trough widths of spike waveforms (MATLAB function *fitgmdist*). We used the intersection of the two gaussian fits as the threshold for separating narrow and wide waveform neurons. Waveforms were obtained by averaging across all recorded spikes for each neuron. Separate models were used for the M1 and striatal populations. We note that we found a larger percentage of narrow waveform neurons that have been previously reported in the literature for the striatum (Extended Data Fig. 5b)^79,113^. However, this high percentage is likely due to the high firing rate of these neurons relative to wide-waveform neurons, especially when looking across a large range of behaviors (Extended Data Fig. 5c), which would make these units more readily detectable with our recording methods.

### Behavior state maps

We adapted the method of Berman et al^43^ that used video image input features to obtain a two-dimensional representation of behavior space, here using muscle activity features instead. We computed behavior state maps for each mouse individually. We first computed spectrograms of each EMG channel using the Morlet wavelet transform, taking 50 frequencies between 0.5 and 20 Hz and using a ω parameter of 5. Channels that did not exhibit clear muscle activity were excluded. This time-frequency transformation was essential for obtaining good segregation of behaviors into clusters, as time-varying muscle activity patterns reflect the idiosyncrasies of each behavior. We embedded the resulting frequency features at each 1 ms time point into a two dimensional space using a MATLAB implementation of the UMAP algorithm^114^ (www.mathworks.com/matlabcentral/fileexchange/71902). We set the number of nearest neighbors to 100, and used a distance metric of either the Euclidean distance, or Kullback–Leibler divergence depending on the mouse, based on which metric was able to better generate distinguishable behavior class clusters. To achieve optimal clustering of behaviors, we computed the UMAP embedding using only the time points that were annotated with one of the ten behavior classes. Additionally, to both avoid the clustering being dominated by short temporal features and reduce the computation time to feasible levels, we only used every 50^th^ time point when computing the embedding.

To generate density plots (Fig. 1e), we divided the UMAP space into 1001 x 1001 equally spaced bins and calculated the fraction of time points within each bin, then smoothed the result by convolving with a two-dimensional gaussian with a 0.2 bin standard deviation. We next segmented the density maps into different regions using the watershed algorithm. To avoid over-segmentation, we first applied the H-minima transform (MATLAB function *imhmin*) on the density plots using a depth value ranging between 0.05-0.20. After implementing the watershed algorithm, we combined the resulting regions into seven main behavioral regions based on the predominant behavior labels of the timepoints within that region. Finally, to assign a behavioral region to all time points in the session, including those not used in the initial UMAP calculation, we projected all time points into the two-dimensional space using the computed UMAP embedding and labeled them based on which behavioral region they fell into.

To visualize the activity of neurons with respect to this behavior space (Fig. 2a), we computed the average firing rate for all the time points lying within in each of the 1001 x 1001 UMAP bins. We then smoothed the firing rates across space by convolution with a two-dimensional gaussian with a 0.3 bin standard deviation.

### Muscle activity during cortical inactivation

To determine the effect of CFA optogenetic inactivation on muscle activity, control events were first identified by shifting each light onset time randomly either 600 ms forward or backwards. We then examined muscle activity in time windows from 100 ms before to 200 ms after light or control event onsets. Before extracting muscle activity segments from each time window, we normalized each muscle recording by dividing by its standard deviation across the whole recording.

For each light or control event, we baseline centered the activity by subtracting the mean value in the 100 ms before onset. We then averaged across all light or all control events to obtain the overall muscle response across all behaviors (Extended Data Fig. 3a). To measure the response for each individual behavioral region, we only used the light onset events that fell within that region. Given the short duration of behavioral events, the plurality of time points from the matching control time series must also occur in that same region. The difference from controls was computed by subtracting each light event time series from the corresponding control event time series. The short latency and long latency effect sizes were calculated by averaging the difference from controls over the first 35 ms or the first 100 ms after light onset, respectively. The reported values (Extended Data Fig. 3d) are averages across all four contralateral forelimb muscles and all three mice.

### Single neuron activity characterization

For each neuron, we calculated the average firing rate in each of the seven behavioral regions by averaging across all time points that fell within that region. To determine if a neuron was modulated within a behavioral region, we generated a distribution of control firing rates by circularly shifting the behavioral region labels in time relative to the neural activity by random amounts (minimum 10 s, and maximum 10 s less than the recording duration) and recalculating the average firing rates for each shift. Neurons were considered modulated for a behavior if their average firing rate during that behavior was at least three standard deviations above the mean of the control firing rate distribution. Neurons were classified as single behavior-selective if they were modulated for only one behavior.

We behavior-normalized the firing rates of each neuron by dividing the mean firing rate in each behavioral region by the sum of firing rates across all regions. This ensures that the mean firing rates across all regions sum to one, facilitating comparison of how uniform or skewed the distributions are for different neurons. To quantify this uniformity, we defined a behavior variability metric, *ρ*, using the following equation:

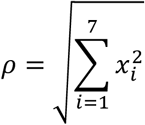

Where 𝑥_𝑖_ is the behavior-normalized average firing rate in the 𝑖th behavioral region. This metric essentially takes the L2 vector norm of the behavior-normalized firing rates, resulting in low values when all the firing rates are uniform and large values when the activity is concentrated within a few behaviors. The minimum value for this metric is 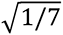, corresponding to the exact same firing rate across all behaviors, and the maximum value is 1, corresponding to the case where only one behavioral region contains non-zero firing rates.

Because the above metric is dependent on how we divided the behavioral regions, we sought to additionally characterize the specificity of neural firing rates in a manner agnostic of the behavior segmentation. The sparsity index has been previously used to characterize place cells in the hippocampus^48,49^. Roughly speaking, it is related to the fraction of space that a cell’s spiking activity occupies. Here, we applied this metric to the activity of M1 and striatal neurons across behavior space rather than physical space. We divided the behavior space into 100 x 100 equally spaced bins and computed the sparsity index as:

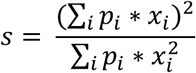

Where 𝑝_𝑖_ is the fraction of the time that the mouse spends in the 𝑖th bin and 𝑥_𝑖_ is the average unnormalized firing rate of the neuron in that bin. We did not perform any spatial smoothing of firing rates for sparsity calculations. To ensure accurate calculation of 𝑝_𝑖_ and 𝑥_𝑖_, we only included bins that contained at least 100 time points.

For both behavior variability and sparsity calculations, we used 10 ms binned firing rates and excluded extremely low firing neurons (mean firing rate < 0.2 spk/s). For muscles, we replaced firing rates with muscle activity levels. Control distributions were obtained by calculating the behavior variability or sparsity after 100 random circular shifts of the behavioral region labels relative to the neural activity, as above.

### Behavior classification analysis

We used a random forest classifier (MATLAB class *TreeBagger*) to determine how well the behavioral region the animal was in could be identified from the neural activity in M1 or striatum. The classifier used 100 decision trees, and used 100 ms binned firing rates with 1 s of history (11 total bins) as input features. To balance class sizes, we randomly subsampled the number of time points in each behavioral region to match the number of time points in the behavioral region with the fewest. To predict future behavior, we shifted the behavior labels forward in time relative to the neural activity in 200 ms increments, up to 4 s into the future. All classification analyses were carried out under 10-fold cross validation and accuracy was reported as the number of correctly labeled time points divided by the total number of time points. We generated 100 different control classifiers by circularly shifting the behavioral region labels a random amount relative to the neural activity in time, as above.

### Comparison of eating and licking activity

To investigate whether the response in eating-specific neurons was primarily related to reward signaling, we leveraged the time points where mice were licking for water. This behavior presumably invokes reward-related activity given the water restriction, but it does not involve movements of the limbs. However, due to the capacitive touch sensors on the water ports, neural recordings during licking from these ports were saturated with electrical artifacts, and unusable for measuring firing rates. Nevertheless, there were a few periods during the experimental session where water dripped down from a port onto the floor or wall, or where we put water droplets on the wall to aid mice in finding the ports for the first time, that we could use to compute firing rates during licking.

We tested whether the eating-specific neurons had similar firing rates during licking as during eating. However, the usable licking time periods were very short and infrequent (9.65 ± 5.87 s of licking in total compared to 328.62 ± 84.79 s of eating, mean ± stdev). To address this, we took 2000 random subsamples of the eating time points to match the number of licking time points, and computed the average firing rate from these subsamples. This allowed us to account for the variance in firing rate calculations when a very small subset of time points were used. After obtaining this distribution of average firing rates during eating, we determined where the average licking firing rate fell among this empirical distribution of eating firing rates, and computed a p-value for each neuron as (𝑘 + 1)⁄(𝑛 + 1) where *k* is the number of eating subsamples with average firing rate less than or equal to the average firing rate during licking, and *n* is the total number of subsamples (n = 2000 here). To account for multiple comparisons, we used the Benjamini-Hochberg correction with a false discovery rate of 0.05. The number of neurons yielding nonsignificant p-values was used to determine the fraction of eating-specific neurons that were similarly responsive during licking (Extended Data Fig. 6f).

### Correlation of average firing rates across behaviors

To examine neural activity similarity across different behaviors or behavioral regions, we took an exhaustive pair-wise approach. For each pair of behaviors being compared, we computed the Spearman correlation of the average firing rates during each of the behaviors across all neurons. From these values, we generated a similarity matrix by using the Spearman correlation across all possible pairs of behaviors (Fig. 3b,g).

We generated a hierarchical cluster tree from this similarity matrix by using single-linkage clustering, with 1 - correlation being the distance metric (MATLAB function *linkage*). Single linkage clustering progressively groups pairs of behaviors or behavior clusters together based on the smallest distance between all pairs. Therefore, behaviors linked together at the low levels of the hierarchy exhibit higher Spearman correlation while behaviors at higher levels exhibit lower Spearman correlation. We visualized the cluster trees using dendrogram plots (MATLAB function *dendrogram*). To quantify the variability of the correlation values across behaviors, we used a hierarchy spread metric, which we defined as the standard deviation in the Spearman correlations across all behavior pairs. To determine the chance level variability in the correlations, we randomly shifted the behavior labels in time relative to the neural activity 100 times, recomputing the Spearman correlations and the hierarchy spread metric each time.

### PCA divergence

We additionally examined the similarity in population covariance structures across all pairs of behaviors or behavioral regions. For each brain area, we performed principal component analysis (PCA) on each of the behaviors separately. That is, for each behavior, we used PCA (MATLAB function *pca*) on 𝑋, the 𝑡 x 𝑛 matrix of 10 ms binned mean-subtracted firing rates, where 𝑡 is the number of time bins used for the given behavior, and 𝑛 is the total number of neurons in M1 or in striatum. To keep the number of time points, 𝑡, the same in the PCA calculations, we subsampled the time points within each behavior (region) to equal the smallest number of time points in any one behavior (region). To mitigate matrix rank-deficiency issues, we excluded extremely low firing rate neurons (mean firing rate <0.2 spk/s). Each PCA calculation produced a matrix of projection weights, 𝑃, corresponding to the principal component (PC) vectors.

Previous studies^13,57^ have quantified the alignment of two different neural activity subspaces using the following method. First, the matrix of neural activity during one behavior is projected into the top 𝑑 PCs computed for that behavior, and the total variance of those projections is computed. Then, the same matrix of neural activity is projected into the top 𝑑 PCs of the second behavior, and the total variance of those projections is computed. The alignment index is then defined as the ratio of those two variances. This can be compactly formulated as:

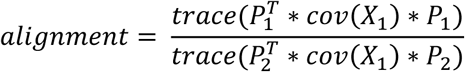

where 𝑋_1_ is the matrix of mean-subtracted neural activity during behavior one, 𝑃_1_ is the matrix of the top 𝑑 PC vectors computed for behavior one, and 𝑃_2_ is the matrix of the top 𝑑 PC vectors computed for behavior two. The value of 𝑑 has been arbitrarily chosen in previous studies with values ranging from 4 to 10^13,57^, but for the analyses in this study, we determined 𝑑 based on the estimate of the dimensionality of the data using parallel analysis^115^ (100 shuffles, alpha = 0.05, using a MATLAB implantation of the algorithm from https://doi.org/10.6084/m9.figshare.13335341).

This alignment index measures how well aligned two subspaces are because projecting the same data into the top PCs of two different population activity measurements should result in similar variance capture if those two measurements exhibit similar covariation across neurons. However, the above metric only takes into account a subset of the PCs and requires choosing a cutoff value 𝑑. Additionally, this metric would misleadingly result in high alignment even if there is drastic changes in the distribution of variance explained in the top 𝑑 PCs since it does not distinguish between individual PCs (for example even if PC 1 and PC 2 completely switched the amount of variance explained in each of them, this metric could still result in high alignment as long as 𝑑 > 2). Here, we introduce an extension of this approach by defining the divergence metric for activity during two behaviors:

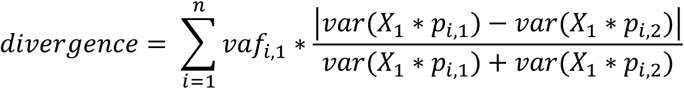

Where 𝑛 is total number of PCs (which equals the total number of neurons), 𝑣𝑎*f*_𝑖,1_ is the fraction of the variance accounted for in the 𝑖th PC in behavior one, 𝑋_1_ is the matrix of mean subtracted firing rates during behavior one, and 𝑝_𝑖,1_ and 𝑝_𝑖,2_ are the projection weights of the 𝑖th PC for behaviors one and two, respectively. 𝑣𝑎*r*(𝑥) indicates variance of 𝑥. Intuitively, for each individual PC, we are taking the absolute difference over the sum of the variances when the same data is projected into that PC for behavior one and projected into that PC for behavior two. We then take a weighted average over all PCs, weighted by the fraction of variance accounted for by the PC in behavior one. Like the alignment index, this divergence metric reflects the similarity of variance captured across different PC projections, but the divergence takes into account variance throughout the full neural activity space, not just a subspace defined by a subset of PCs. Although the later PCs contain low variance when the original data is projected into them, we found that projecting data from other behaviors resulted in higher variance in those PCs (Fig. 3f, Extended Data Fig. 7b). This suggests that the later PCs are not just poorly estimated noise dimensions. Neither the alignment nor the divergence equations are symmetric, so for both, we recomputed the values after switching behaviors one and two and used the mean of the two values.

We also used the first (smallest) principal angle as an additional metric to measure subspace similarity. For the two behaviors, we performed singular value decomposition on the product between their projection matrices of the top 𝑑 PCs, 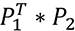. Like with the alignment index, 𝑑 was calculated using parallel analysis. The principal angles were then computed as the inverse cosine of the resulting singular values^28^.

As a control, we performed random rotations of the PCs for the population activity for each behavior, and recomputed the similarity metrics. Random rotation matrices were generated by sampling weights from a standard normal distribution and then orthonormalizing the matrices. We then generated surrogate data matrices, 𝑋_rot_ by performing PCA on the mean-subtracted firing rates, 𝑋, multiplying the resultant matrix of PC time series by the random rotation matrix, then multiplying the result by the inverse of the projection matrix of PCs (which projects the time series back into the original neural dimensions). The resulting surrogate data matrices were then used in place of the matrix of firing rates for performing PCA and subsequently computing the subspace alignment, divergence, and principal angles.

For generating similarity matrices based on activity covariation, we calculated alignment, principle angle, and divergence for all pairs of behavior. To generate the corresponding dendrograms, we used one minus the alignment, the principle angle, or the divergence as the distance metrics respectively. Hierarchy spread and behavior shift controls were computed the same way as for the firing rate correlation hierarchies. For calculating similarity matrices using the ten manually-annotated behaviors rather than the seven UMAP-segmented behavioral regions, we excluded the Jump Across and Still behavior classes due to the small number of labeled time points within those behaviors.

### Population response at muscle activations

To extract muscle activation events for each of the contralateral forelimb muscles, we used a thresholding approach. Activation events were defined as time points where the normalized muscle activity remained below a quiescence threshold *Tq* for at least 150 ms before increasing above an activation threshold *Ta*. Here we used 0.5 standard deviations for *Tq* and 0.7 standard deviations for *Ta*. Events were obtained for each of the muscles and then combined. We extracted the neural firing rates (1 ms bins) in a time window starting from 100 ms before to 200 ms after each event. The activity in M1 or striatum during these activations was calculated by averaging the firing rate across all neurons and across events. To determine the chance-level response, we circularly shifted the muscle activity in time a random amount relative to the neural activity 100 times.

### Muscle correlations

For population-level correlation, we performed canonical correlation analysis (CCA, MATLAB function *canoncorr*) between the neural activity and the activity of the four muscles recorded in the contralateral forelimb. We used the 𝑛 x 𝑡 matrix of mean-subtracted neural activity, 𝑋, where *n* is the number of neurons in M1 or striatum, and *t* is the total number of time points; and the 𝑎 x 𝑡 matrix of mean-subtracted muscle activity, 𝑀, where 𝑎 is the number of muscles (here 𝑎 = 4). CCA finds projection matrices 𝑃_*neur*_ and 𝑃_*musc*_ that maximizes the correlations over time between the projected activity of 𝑋 ∗ 𝑃_*neur*_ and 𝑀 ∗ 𝑃_*musc*_. The canonical correlations are defined as the 𝑎 correlation values attained from these projections. Values reported for Fig. 4e and Fig. 5d-f were the average across all *m* correlations. We obtained chance-level canonical correlations by circularly shifting the muscle activity in time a random amount relative to the neural activity 100 times.

### Muscle activity decoding

To decode muscle activity from the neural activity, we utilized a Wiener cascade decoder^116^. For each muscle, the decoder first generates predictions of the activity, 𝑦(𝑡), using a linear combination of the neural activity, including activity over a preceding time window:

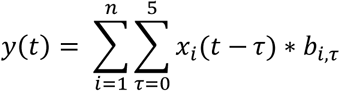

where 𝑛 is the number of neurons, 𝑥_𝑖_(𝑡) is the mean-subtracted firing rate of neuron 𝑖 at time 𝑡, 𝜏 is the bins from each neuron to use as input, and 𝑏_𝑖,𝜏_ is the weight for bin 𝜏 for the given neuron. Here we used 10 ms time bins, and 5 total time bins (i.e. 50 ms) for input. The Weiner cascade decoder also applies a non-linearity after the linear model, here a third-order polynomial:

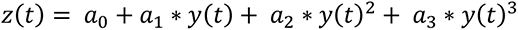

where 𝑎_0_ to 𝑎_3_, are the polynomial coefficients. The linear weights 𝑏_𝑖,𝜏_ are fit from a training dataset of neural activity and muscle activity using least-squares ridge regression (MATLAB function *ridge*, ridge parameter of 1 for striatum, 0.01 for M1). To help prevent matrix rank-deficiency, we excluded extremely low firing rate neurons (mean firing rate < 0.2 spk/s) and we also added a small amount of white noise following a gaussian distribution of mean 0 and standard deviation 0.01 to the decoder inputs. After the linear weights 𝑏_𝑖,𝜏_ were computed, we calculated the estimate 𝑦(𝑡) using those weights and the neural activity, then fit the polynomial weights 𝑎_0_ to 𝑎_3_using least squares polynomial regression (MATLAB function *polyfit*) between the estimated activity 𝑦(𝑡) and the muscle activity.

To measure decoding performance, we defined R^2^ as the multi-channel variance accounted for (mVAF) metric^117^:

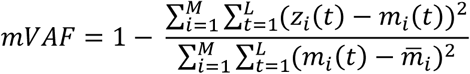

Where 𝐿 is the total number of time points, 𝑀 is the total number of muscles (here 4), 𝑧_𝑖_ (𝑡) is the decoder estimate of muscle activity for muscle 𝑖 at time 𝑡, 𝑎_𝑖_(𝑡) is the actual muscle activity of muscle 𝑖 at time 𝑡, and [inline[is the average muscle activity of muscle 𝑖 across all time points. This is equivalent to taking the weighted average of the traditional VAF for each muscle, with the weight being the fraction of the total variance represented by each muscle. Due to the sensitivity of this metric to extreme outliers, which we occasionally observed when decoding using striatal inputs, we excluded the time points with values beyond the 99th percentile of decoded values in this calculation. We note that this exclusion mainly served to bring up the R^2^ values for striatum decoding from extremely negative values, meaning our results showing better decoding performance in M1 than striatum are made more conservative using this exclusion.

All decoding analyses were performed under 10-fold cross validation. To keep the amount of training data consistent when examining decoding performance during individual behaviors, we subsampled the total number of time points to match the behavior with the fewest time points. Chance-level decoding was assessed by circularly shifting muscle activity in time a random amount relative to neural activity 100 times.

### Muscle correlation stability

To measure how similar the correlations between neurons and muscles were across two behaviors, we computed the Pearson correlation for the activity of every neuron-muscle pair during each pair of behaviors. We then determined whether neurons that were correlated with muscles in one behavior were similarly correlated in the other behavior by taking the Spearman correlation of these neuron-muscle correlations across the two behaviors using all neuron-muscle pairs. We computed this Spearman correlation for every pair of behaviors, and generated dendrograms using one minus the correlation as the distance. The hierarchy spread across behaviors was defined the same way as in the *Correlation of average firing rates across behaviors* section.

To determine the stability of neural-muscle correlations on a population level, we first performed CCA on each behavior individually. CCA finds the projections 𝑃_*neur*_ and 𝑃_*musc*_ that maximize the correlations between the projected neural activity and projected muscle activity. We reasoned that if two behaviors had similar correlations in their neural and muscle activity, then projecting the activity from one behavior into the spaces defined by the projection matrices computed during the second behavior would result in similar canonical correlation values. We therefore measured the similarity in neural-muscle correlations by computing the drop in canonical correlation:

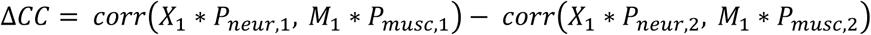

where 𝑋_1_ is the mean-centered matrix of firing rates in behavior one, 𝑀_1_ is the mean-centered matrix of muscle activity in behavior one, 𝑃_*neur*,1_ and 𝑃_*musc*,1_ are the projection matrices for the neural activity and muscle activity, respectively, for behavior one, 𝑃_*neur*,2_ and 𝑃_*musc*,2_ are the same for behavior two, and *corr*(𝑥, 𝑦) denotes the pairwise Pearson correlation between the columns of 𝑥 and 𝑦. Δ𝐶𝐶 is a four-element vector of the drop in the correlations. We then took the average of this drop across all four canonical correlations. Like our divergence metric, this equation is not symmetric, so we recomputed Δ𝐶𝐶 after switching behavior one and behavior two, and took the mean of the two values. We determined chance-level changes in individual neuron-muscle correlations and canonical correlations by circularly shifting the behavior labels a random amount in time relative to the neural and muscle activity 100 times (neural and muscle activity were still aligned in time).

### Recurrent neural network model

To model a muscle activity pattern generating network, we used a recurrently connected artificial neural network (RNN). We used the previously reported PsychRNN package in Python^118^ which we configured to implement a “basic” RNN architecture. Briefly, the model consists of *n* artificial neurons whose activity dynamics are governed by the difference equation:

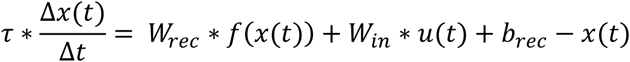

where 𝑥(𝑡) is the vector of activity in the artificial neurons at time 𝑡, 𝑢(𝑡) is the vector of inputs to the network at time 𝑡, 𝑣(𝑥) is a nonlinear transfer function, 𝑊_*rec*_ is the matrix of recurrent network weights, 𝑊_𝑖𝑛_ is the matrix of the input weights, 𝑏_*rec*_ is the vector of bias values for the recurrent units, and 𝜏 is a time constant. The output of the network 𝑧(𝑡) is determined by:

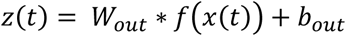

where 𝑊_*out*_ is the matrix of output weights, and 𝑏_*out*_ is the vector of bias values for the outputs. Here we used a rectified linear function as the transfer function, and a 𝜏 value of 1. Our network consisted of 250 recurrent units. It used the mean squared error as the loss function, and did not use regularization. The 𝑊_*rec*_, 𝑊_𝑖𝑛_, 𝑊_*out*_, 𝑏_*rec*_, and 𝑏_*out*_ weights were trained using the backpropagation through time algorithm with Adam optimizer.

We split our data with a ratio of 2:1:1 into train, test, and validation blocks respectively. We used the validation trials to inform when to terminate network training. We trained the network until the validation loss fell below a set threshold, or the validation loss increased by an amount exceeding a threshold (indicating overfitting), or after 150,000 iterations, whichever came first.

We set the outputs of the RNN to be the trial-averaged muscle activity of the four contralateral forelimb muscles. Using the video recording, we manually annotated time points corresponding to one particular movement type during each different behavior. These were: the hand hitting the ground during walking, the hand leaving a handhold to reach for the next handhold during climbing, the arm being raised to bring food to the mouth during eating, the arm being raised to perform a face stroke during grooming, the hand leaving contact with the ground during rearing, and the hand hitting the ground during jumping down from the wall. Because the video recordings had limited temporal resolution, we refined the alignment of the events within each movement category by shifting each annotated event time to maximize muscle activity correlation with the average muscle activity across all events in the given movement category. After alignment, we extracted the neural and muscle activity (in 5 ms time bins) in a time window from 100 ms before to 200 ms after each event.

For the inputs to the network, we used the firing rates of all striatal neurons. We found that the striatal firing rate inputs were unable to drive the network to accurately generate muscle activity on a single-trial basis. We therefore used trial-averaged activity as the inputs and outputs for the network. To reduce overfitting to a single sample of trial-averaged data, on each training iteration we used the trial average of a different random subset of half of the training trials.

To measure how well the network was able to generate muscle activity, we took the Pearson correlation between the actual muscle activity and the network output averaged across the testing trials. As a control, we used constant-value inputs, using a different integer value from 0 to 5 for each of the six movement categories. We also generated control performance values by circularly shifting the striatal activity in time a random amount relative to the muscle activity and repeating the network fitting 100 different times.

## Supplementary materials

Supplementary Video 1. Left, video recording of a mouse performing various behaviors within the arena. Right, plot showing the position in UMAP space at each video frame (open circle). Colored dots denote action potentials for three example neurons.

## Acknowledgments

The authors would like to thank Dr. Andrew Fink and Dr. Lee Miller for their helpful comments on the manuscript. D.X. was supported by the GMCMD T32 training grant at Northwestern University. J.G was supported by R00 NS119787. A.M. was supported by a Searle Scholar Award, a Sloan Research Fellowship, a Whitehall Research Grant Award, The Chicago Biomedical Consortium with support from the Searle Funds at The Chicago Community Trust, the Simons Foundation, and NIH grant DP2 NS120847. Microscopy was performed at the Biological Imaging Facility at Northwestern University (RRID:SCR_017767), graciously supported by the Chemistry for Life Processes Institute, the NU Office for Research, the Department of Molecular Biosciences. This work also made use of the BioCryo facility (RRID:SCR_021288) of Northwestern University’s NUANCE Center, which has received support from the SHyNE Resource (NSF ECCS- 2025633), the IIN, and Northwestern’s MRSEC program (NSF DMR-2308691).

## Author Contributions

The arena was designed and built by D.X. with guidance from A.M. Implantation surgeries were performed by D.X. and A.M. and data collection experiments were carried out by D.X. Data analysis methods were developed and applied by D.X., J.G., and A.M. All authors contributed to the writing and editing of the manuscript.

## Materials & Correspondence

Correspondence and request for materials should be addressed to A.M.

## Data availability

The data that support the findings of this study are available from the corresponding author upon reasonable request.

## Code availability

All Matlab code for data analyses will be made available on GitHub upon publication.

---

**Extended Data Figure 1.**
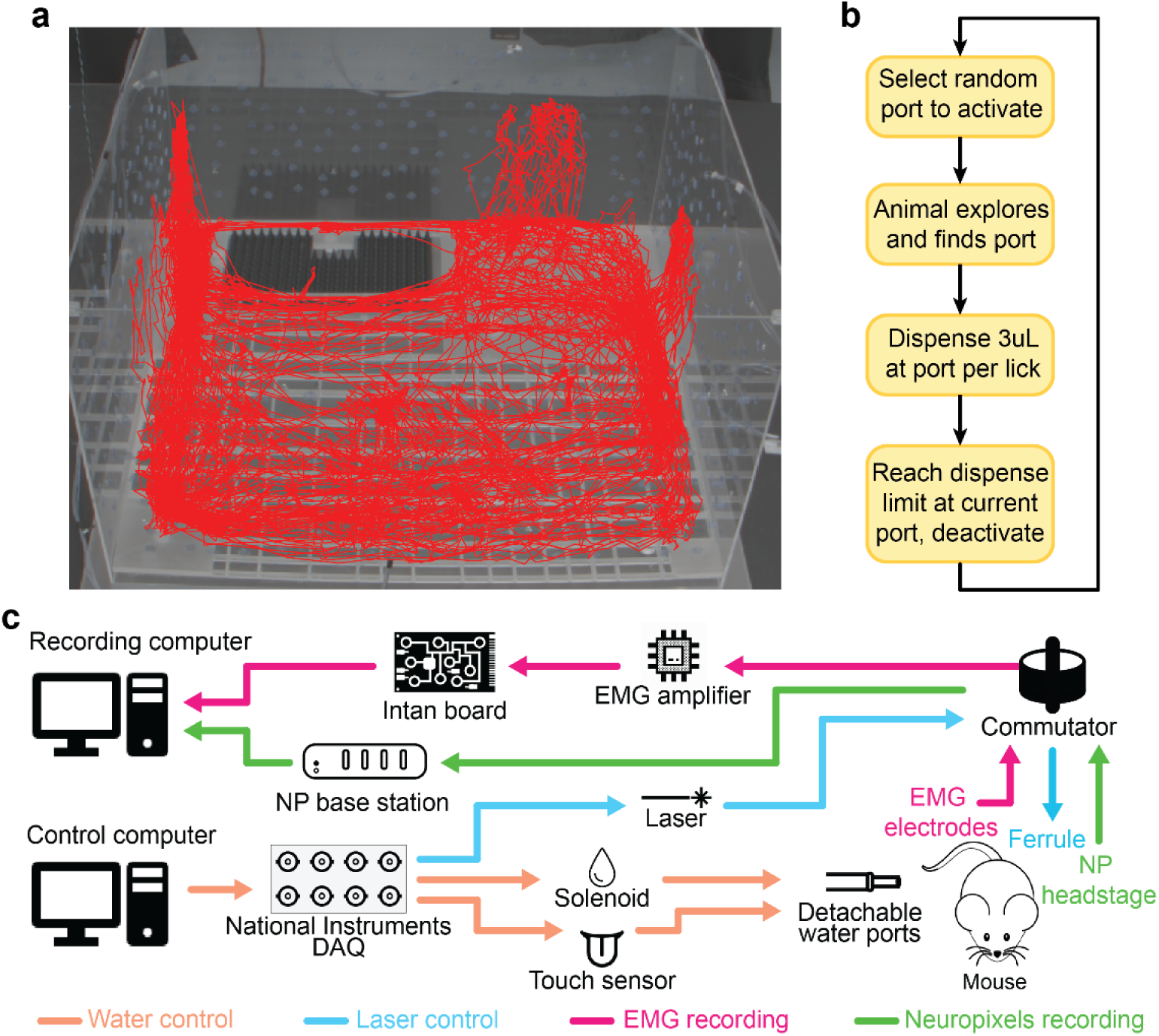
| Paradigm and recording setup. a, Position of one mouse within the arena during one example session. **b,** Water dispensing scheme. **c,** Control and recording system for simultaneous collection of EMG and neural activity. The system also enables control of water ports or laser stimulation for optogenetic inactivation.

**Extended Data Figure 2.**
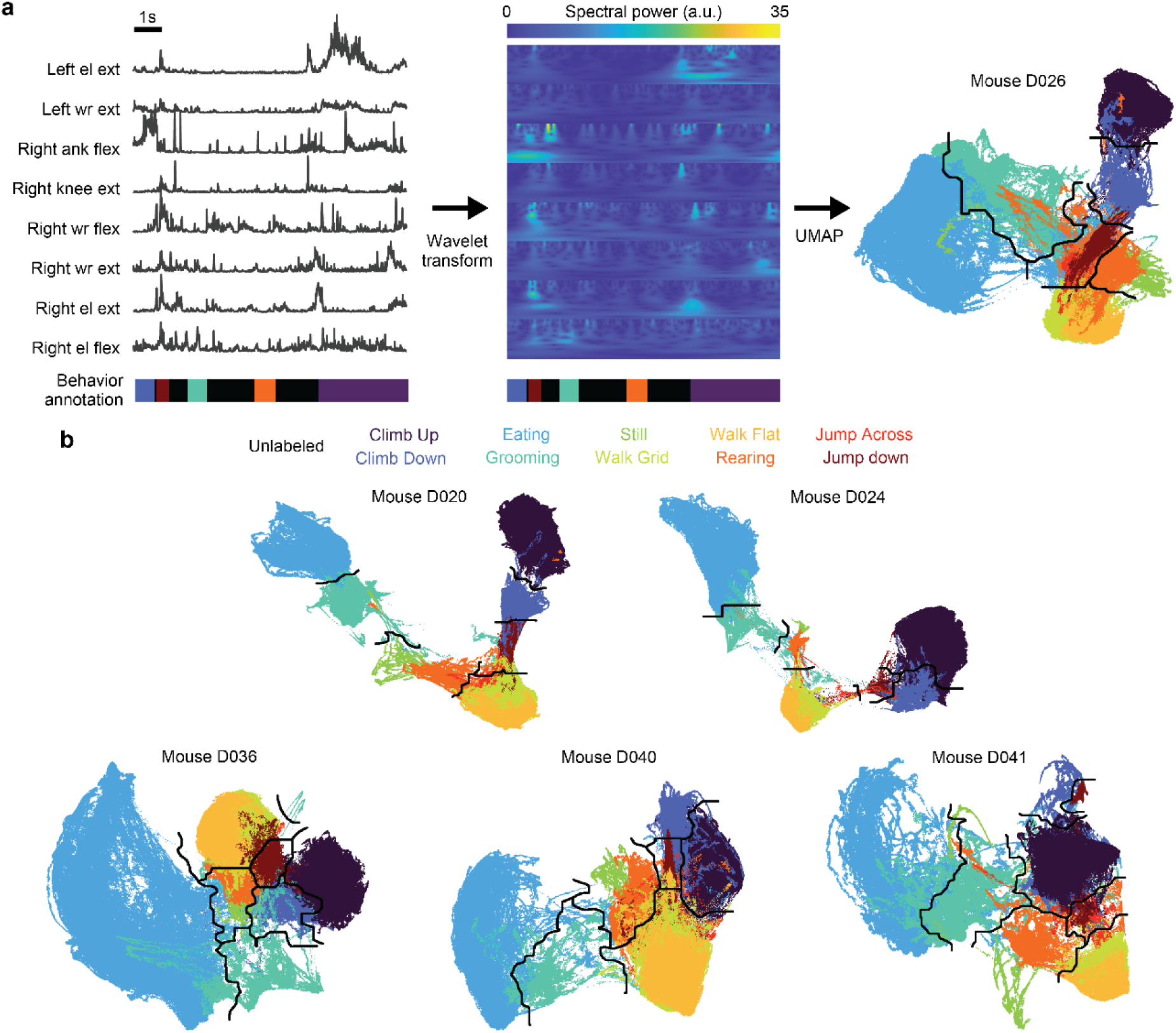
| Construction of the behavior space. a, UMAP processing pipeline. Frequency information is extracted from each of the muscle activity channels using the wavelet transform. These high dimensional frequency features are then projected into a two dimensional space using UMAP. Boundaries between behavioral regions were obtained using the watershed algorithm. UMAP projections and boundaries were computed for each mouse individually. **b,** UMAP behavior spaces for each mouse. Embeddings exhibited well-separated clusters and a similar organization of behavioral regions.

**Extended Data Figure 3.**
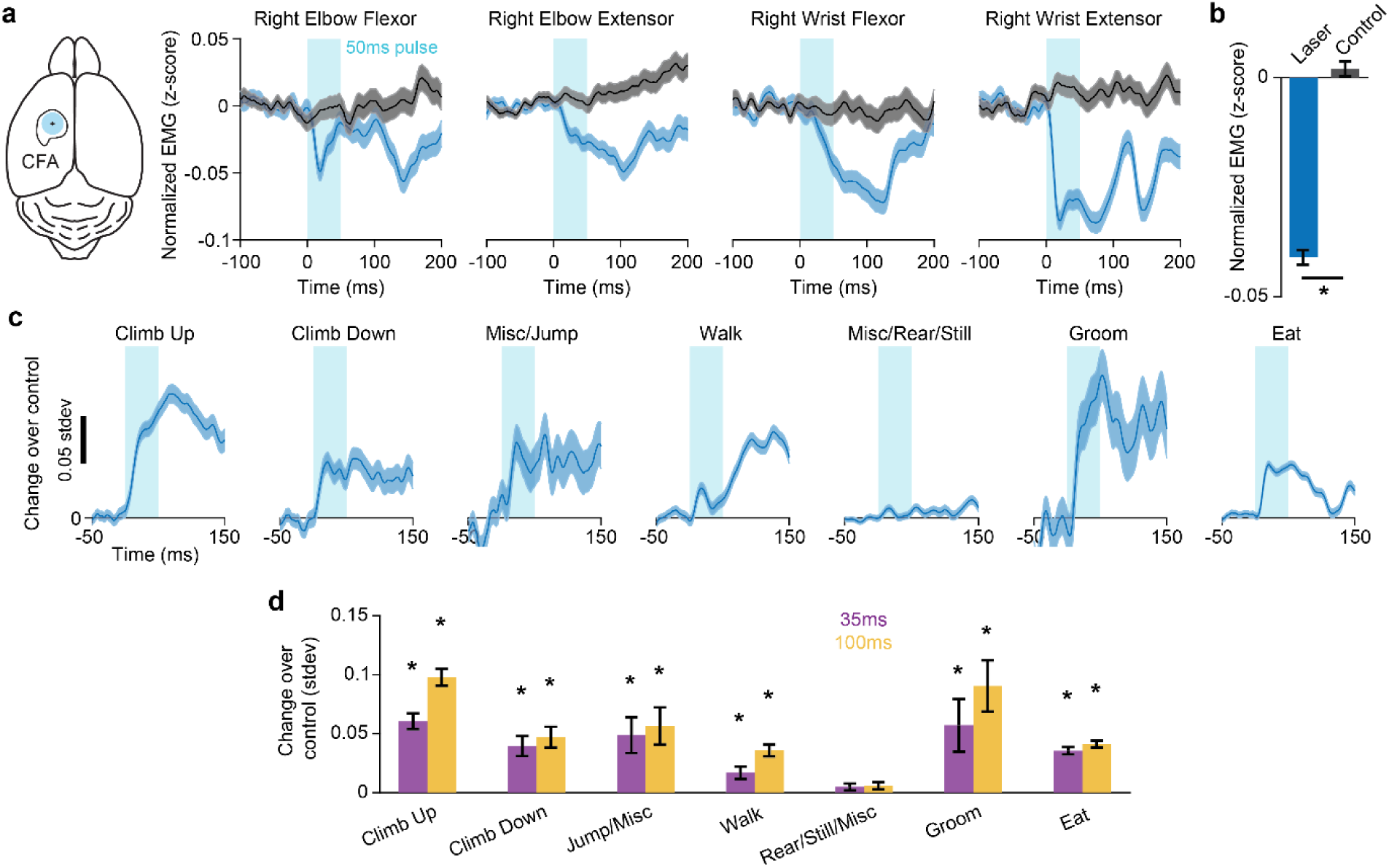
| Motor cortex exerts fast-timescale influence on muscles across most behavioral regions. a, Left, inactivation region. Right, average change in muscle activity in response to optogenetic inactivation of CFA (blue) or no light control (gray) across all four contralateral forelimb muscles. Controls were obtained by sampling from time periods without light inactivation (see Methods). Plots show example session from one mouse, combined across all behaviors; shaded regions denote SEM. **b,** Change in muscle activity between control and inactivation trials combined across all contralateral forelimb muscles of all mice. Two-sided paired t-test, p < 10^-3^. **c,** Difference in muscle activity between control and inactivation trials, averaged across all contralateral forelimb muscles of all mice, for each behavioral region. **d,** Decrease in muscle activity between control and inactivation trials averaged across the first 35 ms or the first 100 ms after light onset. Error bars denote SEM. Stars indicate statistical significance (two-sided paired t-test, with Benjamini-Hochberg multiple comparisons correction, FDR < 0.05)

**Extended Data Figure 4.**
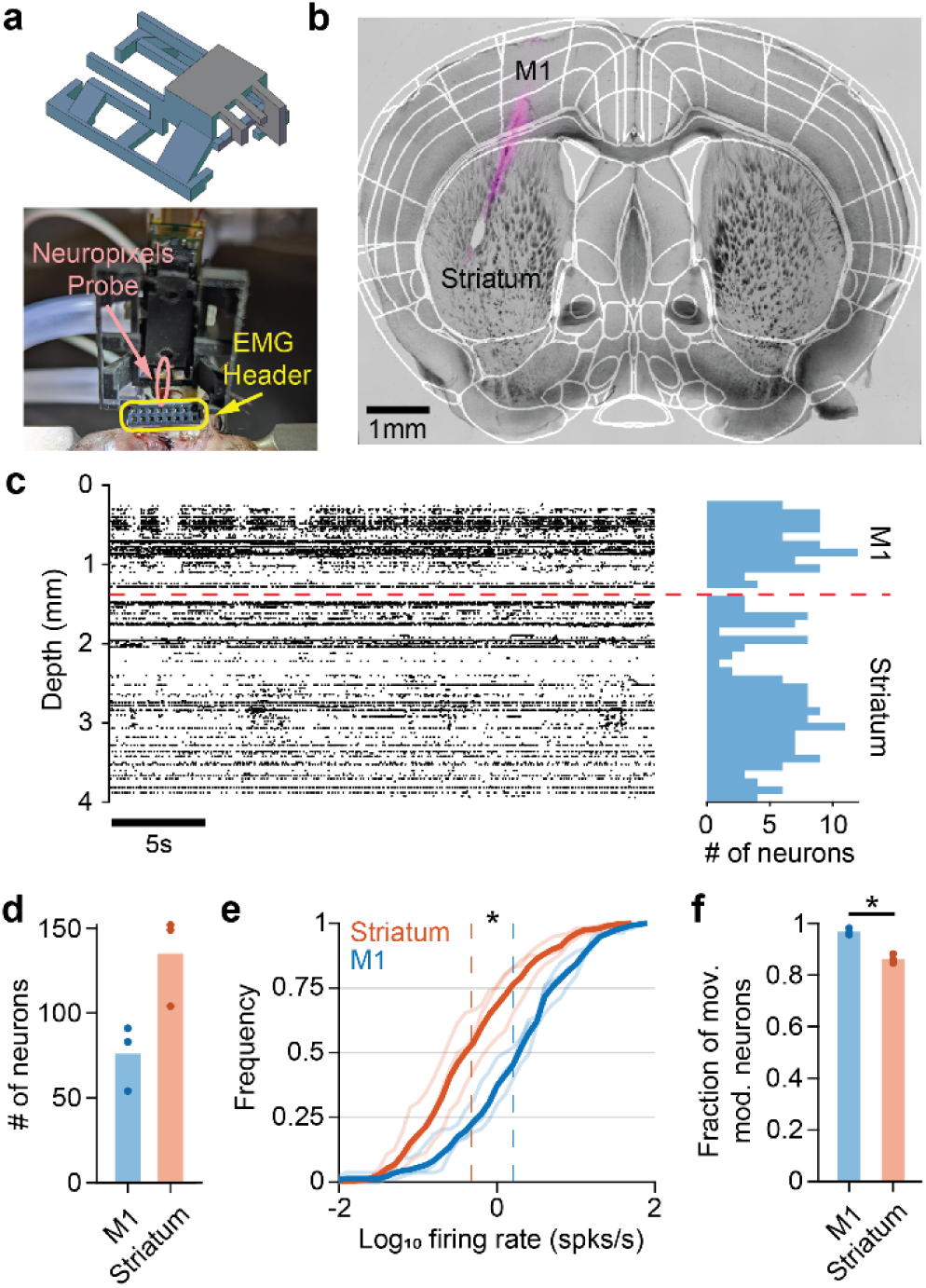
| Chronic neuropixels recordings. a, Top, head fixture design housing the Neuropixels probe and headstage. Modified from Juavinett et al.^50^ Bottom, image of an inserted probe attached to the head of an animal along with a connecter for the EMG electrodes. **b,** Probe insertion location overlayed with the Allen CCF. **c,** Example raster plot of recorded neural activity (left) and the distribution of neurons along the depth of the probe (right). Red dotted line indicates the boundary between M1 and striatum. **d,** Number of neurons recorded in each area. Dots denote individual mice, bars denote averages across mice. **e,** Cumulative histograms of the firing rates averaged across the whole session for neurons recorded in each area. Dotted lines indicate the average across all neurons in each area (two-sided paired t-test, p = 0.0185). **f,** Fraction of neurons in the population modulated by muscle activity above chance (see methods, two-sided paired t-test, p = 0.0269).

**Extended Data Figure 5.**
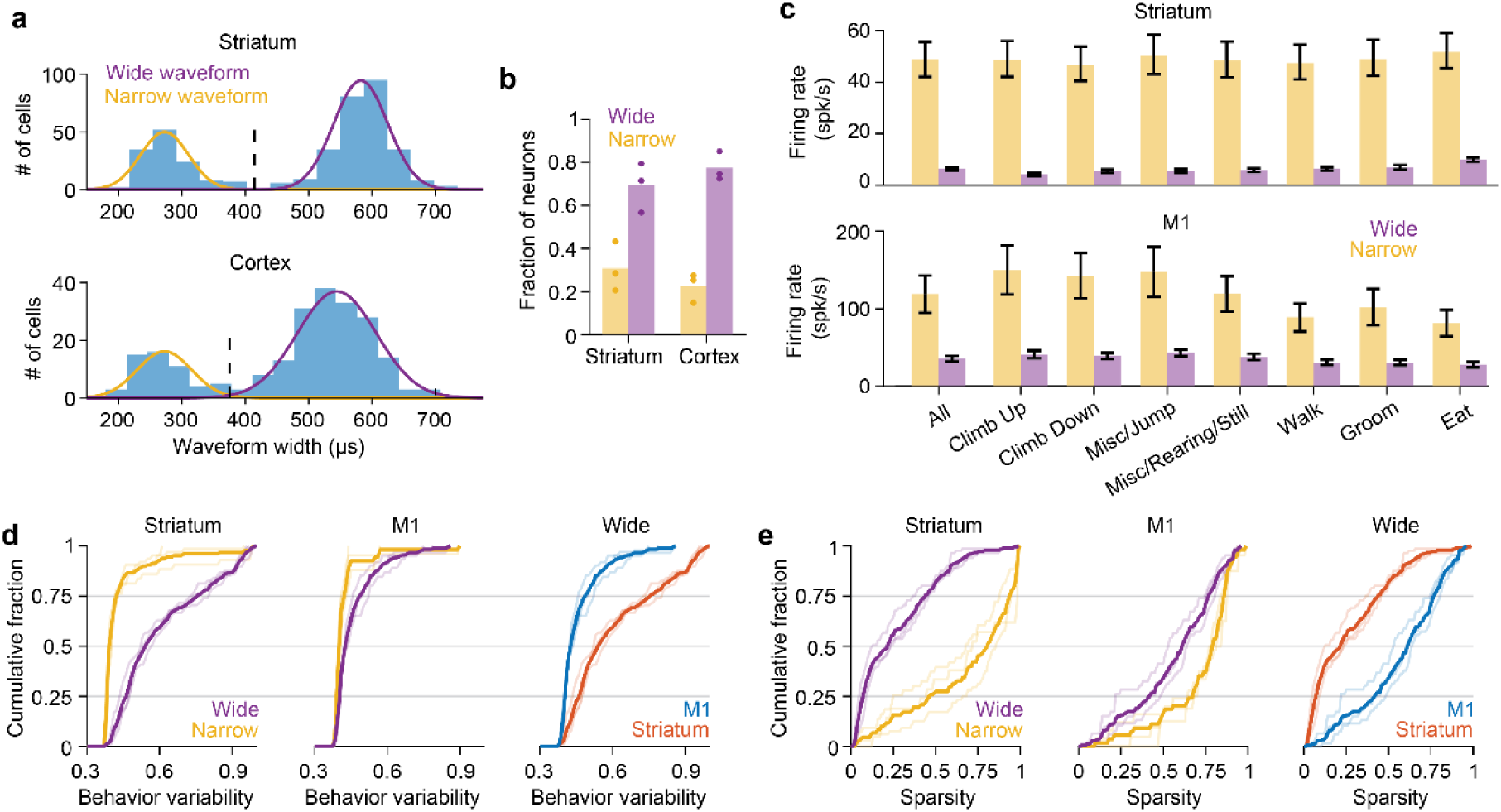
| Distinct properties of wide and narrow-waveform neurons. a, Distribution of spike widths. Curves indicate the gaussian mixture model fit to the distributions. Dotted line indicates the threshold for separating wide and narrow waveforms. **b,** Fraction of each type of neuron in each brain region. **c,** Average firing rate of each type of neuron in each brain region for each behavioral region or all behaviors combined. **d,** Cumulative histograms of behavior variability metrics (*ρ*) for wide and narrow waveform neurons in striatum (left) or M1 (middle) and cumulative histograms of *ρ* for striatal and M1 populations using only wide waveform neurons (right). **e,** Same as in **d**, except for the sparsity index, *s*.

**Extended Data Figure 6.**
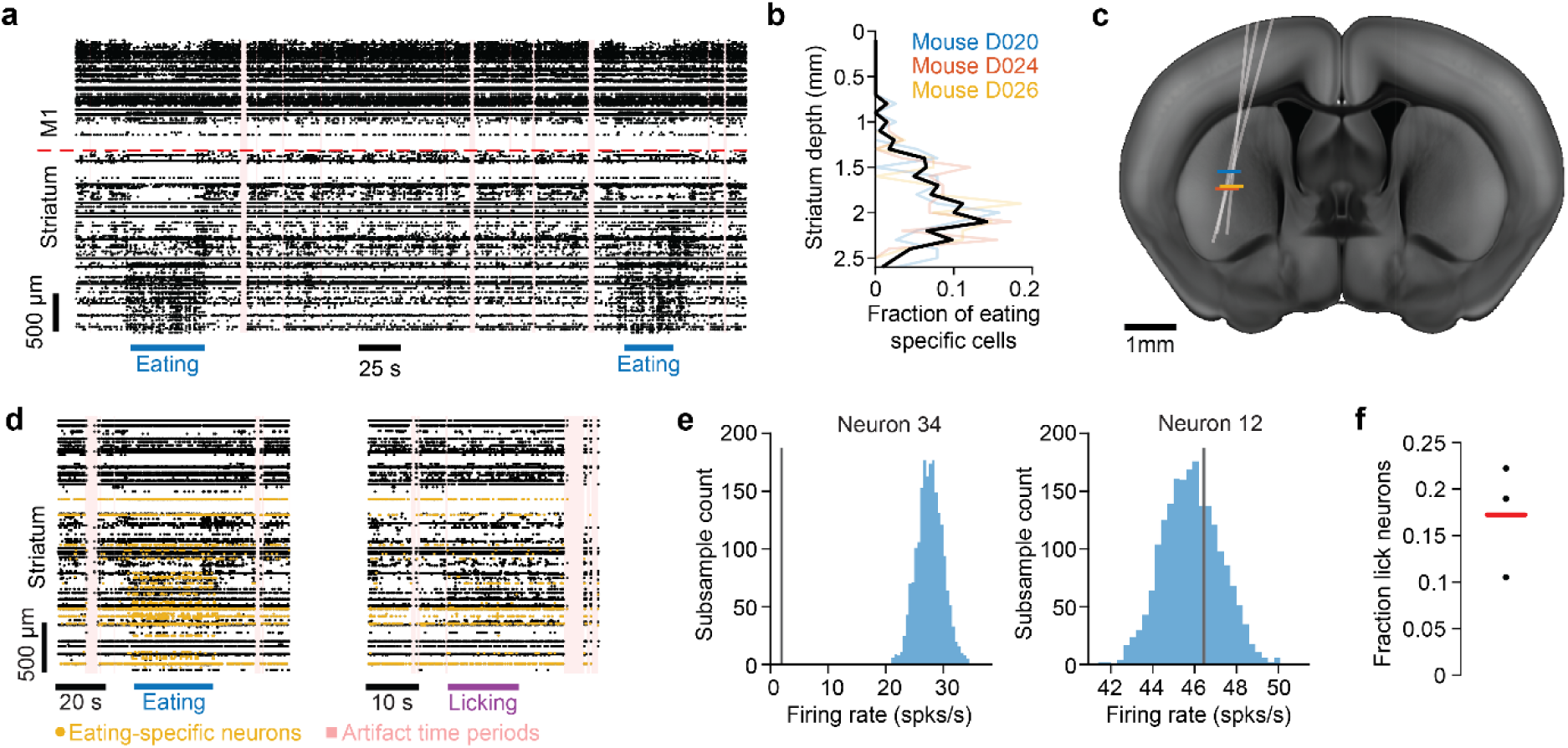
| Spatiotemporal organization of striatal specificity. a, Example raster plot of neurons which includes two bouts of eating. Dotted red line indicates the boundary between M1 and striatum. Pink regions denote time points excluded due to artifacts. **b,** Distribution of eating-specific neurons along the depth of striatum for individual mice (thin, colored) or averaged across mice (thick, black). **c,** Recording probe trajectory for each mouse. Horizontal lines denote boundary within the striatum separating eating-specific neurons (defined as the depth which captures 90% of all eating- specific neurons). **d,** Raster plot showing activity around one eating bout (left) and one licking bout (right) in an example mouse. Yellow denotes eating specific neurons. **e,** Histograms of average firing rates for two example neurons computed for 2000 different random subsamples of eating time points, each matching the number of licking time points (see Methods). Vertical line indicates the average firing rate during licking. Left panel shows a neuron not active during licking (p = 0.0005), right panel shows a neuron active during licking (p = 0.67). **f,** Fraction of eating-specific neurons which are also active during licking. Dots denote individual mice, bars denote averages across mice.

**Extended Data Figure 7.**
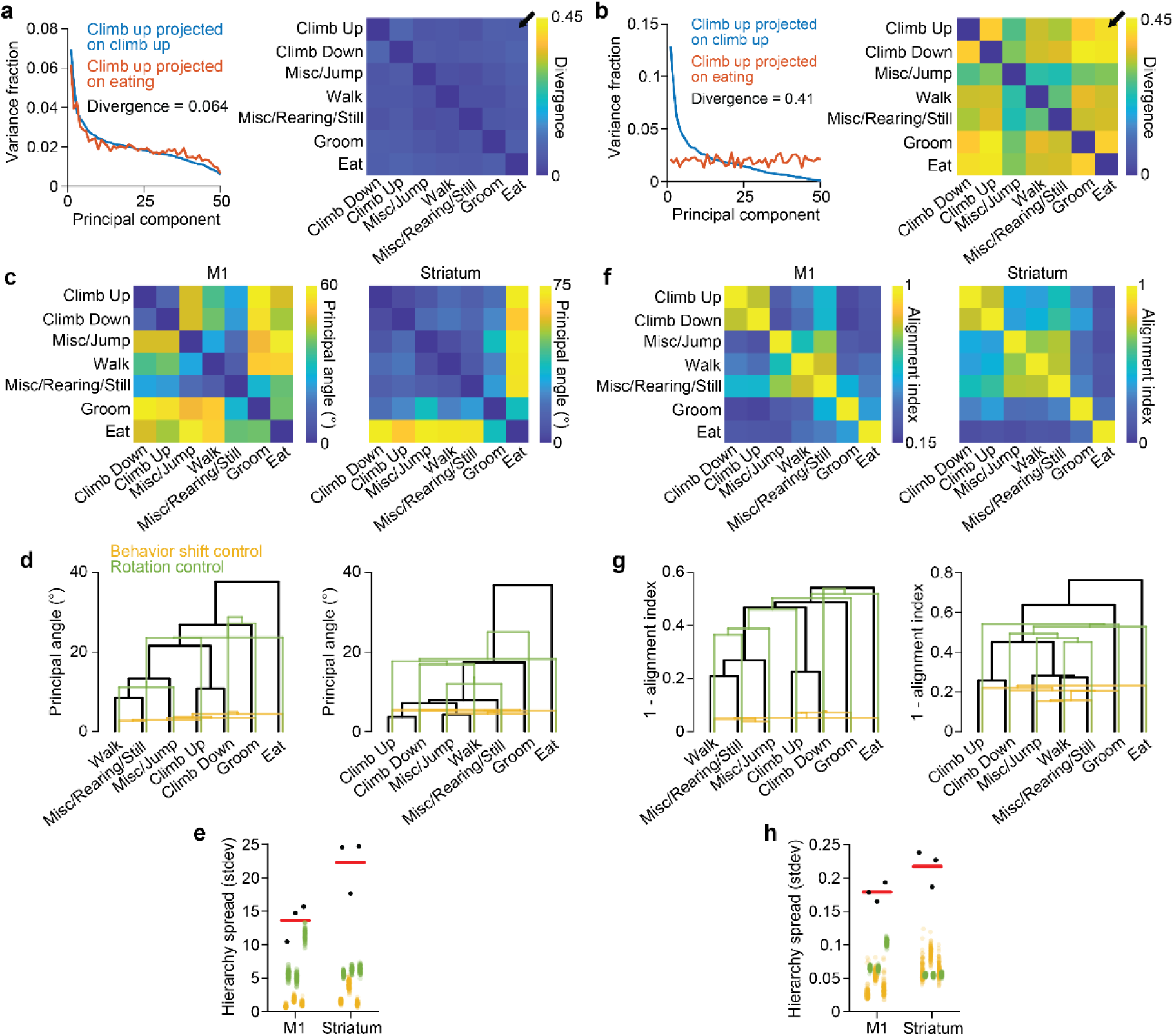
| Covariance divergence controls. a, Left, same as in Figure 3f bottom, except for an example control where behavior labels were randomly shifted in time. Right, the divergence for all pairs of behaviors in the example shift control. Arrow indicates the behavior pair shown in the left panel. **b,** Same as in **a**, except for an example control with random rotations of the PCs. **c,** First principal angle between subspaces for all pairs of behaviors for the same example mouse in Figure 3. **d,** Dendrograms for the first principal angle between behavior subspaces. **e,** Hierarchy spreads for the dendrograms based on first principal angles. **f,g,** Matrices and dendrograms for the subspace alignment index computed for all behavior pairs. **h,** Same as in **e**, except for distances defined as 1 - alignment index.

**Extended Data Figure 8.**
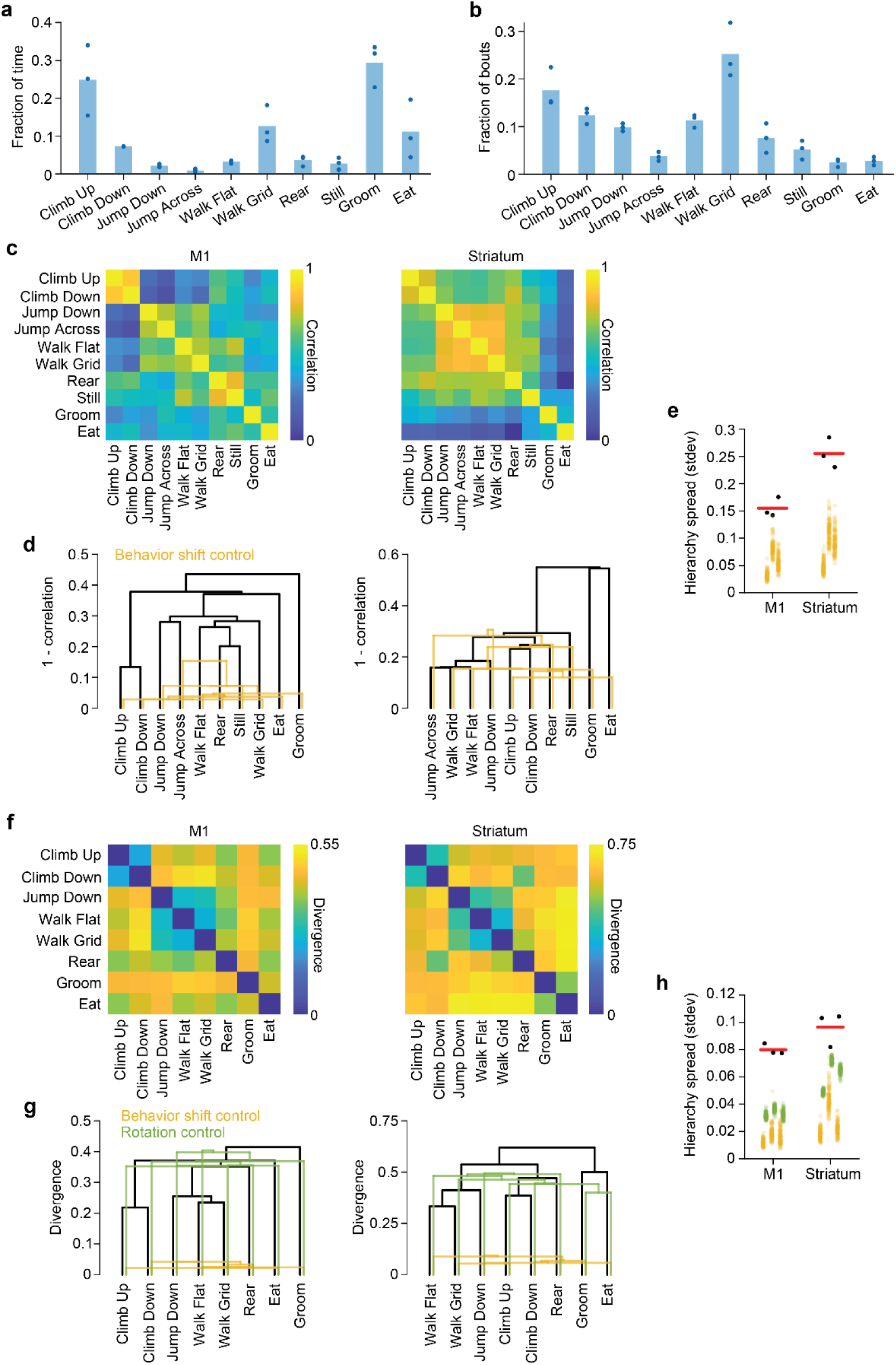
| Hierarchy using epochs based solely on video annotated behaviors. a, Fraction of time points annotated as each of the ten behavior classes. Dots represent individual mice. **b,** Fraction of bouts (continuous epochs) annotated as each behavior class. **c,** Spearman correlation between the average firing rates of individual neurons during pairs of behaviors for one example mouse. **d,** Dendrograms defined using 1 - correlation as the distance. Yellow is one example dendrogram after time shifting the behavior labels. **e,** Hierarchy spread of the dendrograms. For both **e** and **h**, dots indicate individual mice, red bar indicates the average across mice. Yellow dots correspond to 100 behavior shift controls for each mouse. **f,g,** Same as in **c,d** except with divergence. Green indicates one example dendrogram after random rotations of the PCs. **h,** Same as in **d**, except for dendrograms defined using divergence. Green dots denote 100 random rotation controls for each mouse.

**Extended Data Figure 9.**
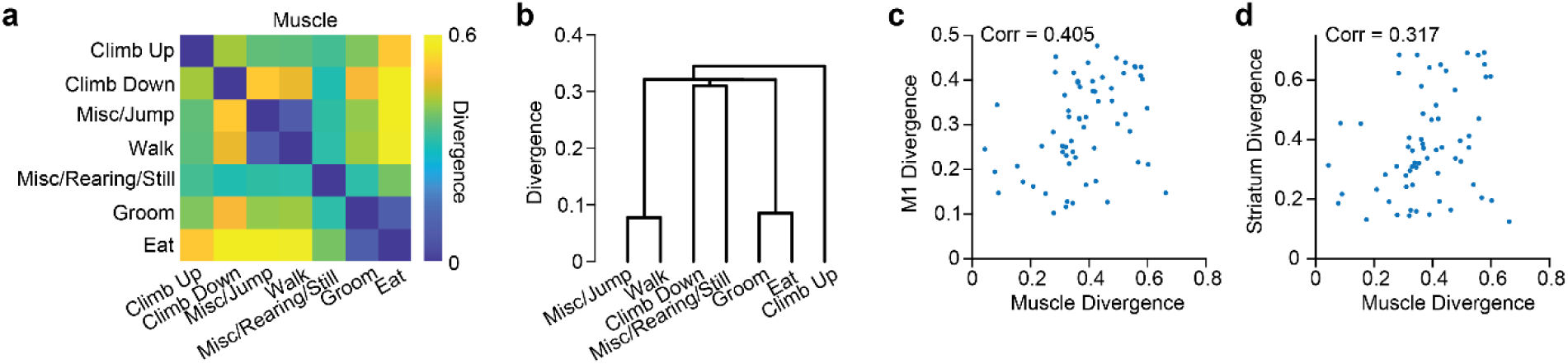
| Muscle activity divergence hierarchy. a, Divergence in muscle activity between all pairs of behavioral regions for one mouse. **b,** Dendrogram for the divergence across behaviors. **c,** The divergence in muscle activity versus the divergence in M1 activity for all behavior pairs combined across all mice. Value in the inset indicates the Spearman correlation. **d,** Same as in **c**, except for striatum rather than M1 divergence.

## References

1. Warriner, C. L., Fageiry, S. K., Carmona, L. M. & Miri, A. Towards Cell and Subtype Resolved Functional Organization: Mouse as a Model for the Cortical Control of Movement. Neuroscience vol. 450 151–160 at 10.1016/j.neuroscience.2020.07.054 (2020).

2. Krakauer, J. W., Ghazanfar, A. A., Gomez-Marin, A., MacIver, M. A. & Poeppel, D. Neuroscience Needs Behavior: Correcting a Reductionist Bias. Neuron vol. 93 480–490 at 10.1016/j.neuron.2016.12.041 (2017).

3. Graziano, M. The organization of behavioral repertoire in motor cortex. Annu. Rev. Neurosci. 29, 105–34 (2006).

4. Cisek, P. & Kalaska, J. F. Neural Mechanisms for Interacting with a World Full of Action Choices. Annu. Rev. Neurosci. 33, 269–298 (2010).

5. Harrison, T. C. & Murphy, T. H. Motor maps and the cortical control of movement. Current Opinion in Neurobiology vol. 24 88–94 at 10.1016/j.conb.2013.08.018 (2014).

6. Dombeck, D. A., Graziano, M. S. & Tank, D. W. Functional clustering of neurons in motor cortex determined by cellular resolution imaging in awake behaving mice. J. Neurosci. 29, 13751–13760 (2009).

7. Yang, W., Kanodia, H. & Arber, S. Structural and functional map for forelimb movement phases between cortex and medulla. Cell 186, 162–177.e18 (2023).

8. Ruder, L. et al. A functional map for diverse forelimb actions within brainstem circuitry. Nature 590, 445–450 (2021).

9. Wang, X. et al. Deconstruction of Corticospinal Circuits for Goal-Directed Motor Skills. Cell 171, 440–455.e14 (2017).

10. Suresh, A. K. et al. Neural population dynamics in motor cortex are different for reach and grasp. Elife 9, 1–16 (2020).

11. Warriner, C. L., Fageiry, S., Saxena, S., Costa, R. M. & Correspondence, A. M. Motor cortical influence relies on task-specific activity covariation. (2022) doi:10.1016/j.celrep.2022.111427.

12. Ma, X., Bodkin, K. L. & Miller, L. E. Population activity in motor cortex is influenced by the contexts of the motor behavior. in International IEEE/EMBS Conference on Neural Engineering, NER vols 2021-May 1152–1155 (IEEE Computer Society, 2021).

13. Miri, A. et al. Behaviorally Selective Engagement of Short-Latency Effector Pathways by Motor Cortex. Neuron 95, 683–696.e11 (2017).

14. Georgopoulos, A. P., Kalaska, J. F., Caminiti, R. & Massey, J. T. On the relations between the direction of two-dimensional arm movements and cell discharge in primate motor cortex. J. Neurosci. 2, 1527–1537 (1982).

15. Schwartz, A. B., Kettner, R. E. & Georgopoulos, A. P. Primate motor cortex and free arm movements to visual targets in three-dimensional space. I. Relations between single cell discharge and direction of movement. J. Neurosci. 8, 2913–2927 (1988).

16. Paninski, L., Fellows, M. R., Hatsopoulos, N. G. & Donoghue, J. P. Spatiotemporal Tuning of Motor Cortical Neurons for Hand Position and Velocity. J. Neurophysiol. 91, 515–532 (2004).

17. Hocherman, S. & Wise, S. P. Effects of hand movement path on motor cortical activity in awake, behaving rhesus monkeys. Exp. brain Res. 83, 285–302 (1991).

18. Saleh, M., Takahashi, K. & Hatsopoulos, N. G. Encoding of Coordinated Reach and Grasp Trajectories in Primary Motor Cortex. J. Neurosci. 32, 1220–1232 (2012).

19. Sergio, L. E. & Kalaska, J. F. Systematic changes in motor cortex cell activity with arm posture during directional isometric force generation. J. Neurophysiol. 89, 212–228 (2003).

20. Scott, S. H. & Kalaska, J. F. Reaching movements with similar hand paths but different arm orientations. I. Activity of individual cells in motor cortex. J. Neurophysiol. 77, 826–852 (1997).

21. Aflalo, T. N. & Graziano, M. S. A. Partial tuning of motor cortex neurons to final posture in a free-moving paradigm. Proc. Natl. Acad. Sci. U. S. A. 103, 2909– 2914 (2006).

22. Cheney, P. D. & Fetz, E. E. Functional classes of primate corticomotoneuronal cells and their relation to active force. J. Neurophysiol. 44, 773–791 (1980).

23. Maier, M. A., Bennett, K. M. B., Hepp-Reymond, M. C. & Lemon, R. N. Contribution of the monkey corticomotoneuronal system to the control of force in precision grip. J. Neurophysiol. 69, 772–785 (1993).

24. Evarts, E. V. Relation of pyramidal tract activity to force exerted during voluntary movement. J. Neurophysiol. 31, 14–27 (1968).

25. Hatsopoulos, N. G., Xu, Q. & Amit, Y. Encoding of movement fragments in the motor cortex. J. Neurosci. 27, 5105–5114 (2007).

26. Athalye, V. R. et al. Invariant neural dynamics drive commands to control different movements. Curr. Biol. 33, 2962–2976.e15 (2023).

27. Churchland, M. M. et al. Neural population dynamics during reaching. Nature 487, 51–56 (2012).

28. Gallego, J. A. et al. Cortical population activity within a preserved neural manifold underlies multiple motor behaviors. Nat. Commun. 9, 4233 (2018).

29. Yang, G. R., Cole, M. W. & Rajan, K. How to study the neural mechanisms of multiple tasks. Current Opinion in Behavioral Sciences vol. 29 134–143 at 10.1016/j.cobeha.2019.07.001 (2019).

30. Markowitz, J. E. et al. The Striatum Organizes 3D Behavior via Moment-to- Moment Action Selection. Cell 174, 44–58.e17 (2018).

31. Klaus, A. et al. The Spatiotemporal Organization of the Striatum Encodes Action Space. Neuron 95, 1171–1180.e7 (2017).

32. Wiltschko, A. B. et al. Mapping Sub-Second Structure in Mouse Behavior. Neuron 88, 1121–1135 (2015).

33. Hsu, A. I. & Yttri, E. A. B-SOiD, an open-source unsupervised algorithm for identification and fast prediction of behaviors. Nat. Commun. 12, 1–13 (2021).

34. Latham, N. & Mason, G. From house mouse to mouse house: the behavioural biology of free-living Mus musculus and its implications in the laboratory. Appl. Anim. Behav. Sci. 86, 261–289 (2004).

35. Bariselli, S., Fobbs, W. C., Creed, M. C. & Kravitz, A. V. A competitive model for striatal action selection. Brain Research vol. 1713 70–79 at 10.1016/j.brainres.2018.10.009 (2019).

36. Mink, J. W. The basal ganglia: Focused selection and inhibition of competing motor programs. Prog. Neurobiol. 50, 381–425 (1996).

37. Kravitz, A. V. et al. Regulation of parkinsonian motor behaviours by optogenetic control of basal ganglia circuitry. Nature 466, 622–626 (2010).

38. Nonomura, S. et al. Monitoring and Updating of Action Selection for Goal-Directed Behavior through the Striatal Direct and Indirect Pathways. Neuron 99, 1302–1314.e5 (2018).

39. Klaus, A., Alves Da Silva, J. & Costa, R. M. What, If, and When to Move: Basal Ganglia Circuits and Self-Paced Action Initiation. Annual Review of Neuroscience vol. 42 459–483 at 10.1146/annurev-neuro-072116-031033 (2019).

40. Gower, J. C. & Ross, G. J. S. Minimum Spanning Trees and Single Linkage Cluster Analysis. Appl. Stat. 18, 54 (1969).

41. Murtagh, F. & Contreras, P. Algorithms for hierarchical clustering: an overview. WIREs Data Min. Knowl. Discov. 2, 86–97 (2012).

42. Akay, T., Tourtellotte, W. G., Arber, S. & Jessell, T. M. Degradation of mouse locomotor pattern in the absence of proprioceptive sensory feedback. Proc. Natl. Acad. Sci. U. S. A. 111, 16877–16882 (2014).

43. Berman, G. J., Choi, D. M., Bialek, W. & Shaevitz, J. W. Mapping the stereotyped behaviour of freely moving fruit flies. J. R. Soc. Interface 11, (2014).

44. Zhao, S. et al. Cell type-specific channelrhodopsin-2 transgenic mice for optogenetic dissection of neural circuitry function. Nat. Methods 8, 745–755 (2011).

45. Guo, Z. V. et al. Flow of cortical activity underlying a tactile decision in mice. Neuron 81, 179–194 (2014).

46. Li, N. et al. Spatiotemporal constraints on optogenetic inactivation in cortical circuits. Elife 8, (2019).

47. Jun, J. J. et al. Fully integrated silicon probes for high-density recording of neural activity. Nat. 2017 5517679 **551**, 232–236 (2017).

48. Shin, J., Lee, H. W., Jin, S. W. & Lee, I. Subtle visual change in a virtual environment induces heterogeneous remapping systematically in CA1, but not CA3. Cell Rep. 41, 111823 (2022).

49. Jung, M. W., Wiener, S. I. & McNaughton, B. L. Comparison of spatial firing characteristics of units in dorsal and ventral hippocampus of the rat. J. Neurosci. 14, 7347–7356 (1994).

50. Juavinett, A. L., Bekheet, G. & Churchland, A. K. Chronically implanted neuropixels probes enable high-yield recordings in freely moving mice. Elife 8, (2019).

51. Tasic, B. et al. Adult mouse cortical cell taxonomy revealed by single cell transcriptomics. Nat. Neurosci. 19, 335–346 (2016).

52. Patel, A. P. et al. Single-cell RNA-seq highlights intratumoral heterogeneity in primary glioblastoma. Science *(80-.).* **344**, 1396–1401 (2014).

53. Gofflot, F. et al. Systematic Gene Expression Mapping Clusters Nuclear Receptors According to Their Function in the Brain. Cell 131, 405–418 (2007).

54. Gorbalenya, A. E. et al. The new scope of virus taxonomy: partitioning the virosphere into 15 hierarchical ranks. Nature Microbiology vol. 5 668–674 at 10.1038/s41564-020-0709-x (2020).

55. Shenoy, K. V., Sahani, M. & Churchland, M. M. Cortical Control of Arm Movements: A Dynamical Systems Perspective. Annu. Rev. Neurosci. 36, 337– 359 (2013).

56. Gallego, J. A., Perich, M. G., Miller, L. E. & Solla, S. A. Neural Manifolds for the Control of Movement. Neuron 94, 978–984 (2017).

57. Elsayed, G. F., Lara, A. H., Kaufman, M. T., Churchland, M. M. & Cunningham, J. P. Reorganization between preparatory and movement population responses in motor cortex. Nat. Commun. 7, 1–15 (2016).

58. Xing, D., Truccolo, W. & Borton, D. A. Emergence of distinct neural subspaces in motor cortical dynamics during volitional adjustments of ongoing locomotion. bioRxiv 2022.04.03.486001 (2022) doi:10.1101/2022.04.03.486001.

59. Voloh, B. et al. Hierarchical action encoding in prefrontal cortex of freely moving macaques. Cell Rep. 42, 113091 (2023).

60. Felleman, D. J. & Van Essen, D. C. Distributed hierarchical processing in the primate cerebral cortex. Cereb. Cortex 1, 1–47 (1991).

61. Saiki-Ishikawa, A., et al. Hierarchy between forelimb premotor and primary motor cortices and its manifestation in their firing patterns. bioRxiv 2023.09.23.559136 (2024) doi:10.1101/2023.09.23.559136.

62. Merel, J., Botvinick, M. & Wayne, G. Hierarchical motor control in mammals and machines. Nature Communications vol. 10 1–12 at 10.1038/s41467-019-13239-6 (2019).

63. Kiritani, T., Wickersham, I. R., Seung, H. S. & Shepherd, G. M. G. Hierarchical connectivity and connection-specific dynamics in the corticospinal-corticostriatal microcircuit in mouse motor cortex. J. Neurosci. 32, 4992–5001 (2012).

64. Arber, S. & Costa, R. M. Networking brainstem and basal ganglia circuits for movement. Nat. Rev. Neurosci. 2022 1–19 (2022) doi:10.1038/s41583-022-00581-w.

65. Sussillo, D., Churchland, M. M., Kaufman, M. T. & Shenoy, K. V. A neural network that finds a naturalistic solution for the production of muscle activity. Nat. Neurosci. 18, 1025–1033 (2015).

66. Vargas-Irwin, C. E. et al. Decoding complete reach and grasp actions from local primary motor cortex populations. J. Neurosci. 30, 9659–9669 (2010).

67. Kaplan, H. S., Salazar Thula, O., Khoss, N. & Zimmer, M. Nested Neuronal Dynamics Orchestrate a Behavioral Hierarchy across Timescales. Neuron 105, 562–576.e9 (2020).

68. Dawkins, R. Hierarchical organisation: A candidate principle for ethology. in Growing points in ethology. (Cambridge U Press, 1976).

69. Driscoll, L. N., Shenoy, K. & Sussillo, D. Flexible multitask computation in recurrent networks utilizes shared dynamical motifs. Nat. Neurosci. 27, 1349– 1363 (2024).

70. Yang, G. R., Joglekar, M. R., Song, H. F., Newsome, W. T. & Wang, X. J. Task representations in neural networks trained to perform many cognitive tasks. Nat. Neurosci. 22, 297–306 (2019).

71. Kaufman, M. T., Churchland, M. M., Ryu, S. I. & Shenoy, K. V. Cortical activity in the null space: Permitting preparation without movement. Nat. Neurosci. 17, 440– 448 (2014).

72. Hennig, J. A. et al. Learning is shaped by abrupt changes in neural engagement. Nat. Neurosci. 24, 727–736 (2021).

73. Steinmetz, N. A., Zatka-Haas, P., Carandini, M. & Harris, K. D. Distributed coding of choice, action and engagement across the mouse brain. Nature 576, 266–273 (2019).

74. Sauerbrei, B. A. et al. Cortical pattern generation during dexterous movement is input-driven. Nat. 2019 5777790 **577**, 386–391 (2019).

75. Pruszynski, J. A. et al. Primary motor cortex underlies multi-joint integration for fast feedback control. Nature 478, 387–390 (2011).

76. Marino, P. J. et al. A posture subspace in primary motor cortex. bioRxiv 2024.08.12.607361 (2024) doi:10.1101/2024.08.12.607361.

77. Xing, D., Truccolo, W. & Borton, D. A. Emergence of Distinct Neural Subspaces in Motor Cortical Dynamics during Volitional Adjustments of Ongoing Locomotion. J. Neurosci. 42, 9142–9157 (2022).

78. Kirk, E. A., Hope, K. T., Sober, S. J. & Sauerbrei, B. A. An output-null signature of inertial load in motor cortex. Nat. Commun. 15, 1–20 (2024).

79. Minkowicz, S. et al. Striatal ensemble activity in an innate naturalistic behavior. Elife 12, (2023).

80. Parker, J. G. et al. Diametric neural ensemble dynamics in parkinsonian and dyskinetic states. Nature 557, 177–182 (2018).

81. Barbera, G. et al. Spatially Compact Neural Clusters in the Dorsal Striatum Encode Locomotion Relevant Information. Neuron 92, 202–213 (2016).

82. Hintiryan, H. et al. The mouse cortico-striatal projectome. Nat. Neurosci. 2016 198 **19**, 1100–1114 (2016).

83. Lee, J. & Sabatini, B. L. Striatal indirect pathway mediates exploration via collicular competition. Nature 599, 645–649 (2021).

84. Schultz, W., Apicella, P., Scarnati, E. & Ljungberg, T. Neuronal activity in monkey ventral striatum related to the expectation of reward. J. Neurosci. 12, 4595–4610 (1992).

85. O’Doherty, J. et al. Dissociable Roles of Ventral and Dorsal Striatum in Instrumental Conditioning. Science (80-.). 304, 452–454 (2004).

86. Gendelis, S., Inbar, D. & Kupchik, Y. M. The role of the nucleus accumbens and ventral pallidum in feeding and obesity. Progress in Neuro-Psychopharmacology and Biological Psychiatry vol. 111 110394 at 10.1016/j.pnpbp.2021.110394 (2021).

87. O’Connor, E. C. et al. Accumbal D1R Neurons Projecting to Lateral Hypothalamus Authorize Feeding. Neuron 88, 553–564 (2015).

88. Matikainen-Ankney, B. A. et al. Nucleus Accumbens D1 Receptor–Expressing Spiny Projection Neurons Control Food Motivation and Obesity. Biol. Psychiatry 93, 512–523 (2023).

89. Barrett, J. M., Martin, M. E. & Shepherd, G. M. G. Manipulation-specific cortical activity as mice handle food. Curr. Biol. 32, 4842–4853.e6 (2022).

90. An, X., et al. A cortical circuit for orchestrating oromanual food manipulation. bioRxiv 2022.12.03.518964 (2024) doi:10.1101/2022.12.03.518964.

91. Peters, A. J., Fabre, J. M. J., Steinmetz, N. A., Harris, K. D. & Carandini, M. Striatal activity topographically reflects cortical activity. Nature 591, 420–425 (2021).

92. Shepherd, G. M. G. Corticostriatal connectivity and its role in disease. Nat. Rev. Neurosci. 2013 144 **14**, 278–291 (2013).

93. Nelson, A., Abdelmesih, B. & Costa, R. M. Corticospinal populations broadcast complex motor signals to coordinated spinal and striatal circuits. Nat. Neurosci. 2021 2412 24, 1721–1732 (2021).

94. Li, N., Chen, T. W., Guo, Z. V., Gerfen, C. R. & Svoboda, K. A motor cortex circuit for motor planning and movement. Nature 519, 51–56 (2015).

95. Cheney, P. D. & Fetz, E. E. Comparable patterns of muscle facilitation evoked by individual corticomotoneuronal (CM) cells and by single intracortical microstimuli in primates: Evidence for functional groups of CM cells. J. Neurophysiol. 53, 786– 804 (1985).

96. Dhawale, A. K., Wolff, S. B. E., Ko, R. & Ölveczky, B. P. The basal ganglia control the detailed kinematics of learned motor skills. Nat. Neurosci. 2021 249 **24**, 1256–1269 (2021).

97. Desmurget, M. & Turner, R. S. Motor Sequences and the Basal Ganglia: Kinematics, Not Habits. J. Neurosci. 30, 7685 (2010).

98. Rueda-Orozco, P. E. & Robbe, D. The striatum multiplexes contextual and kinematic information to constrain motor habits execution. Nat. Neurosci. 18, 453– 462 (2015).

99. Yttri, E. A. & Dudman, J. T. Opponent and bidirectional control of movement velocity in the basal ganglia. Nat. 2016 5337603 **533**, 402–406 (2016).

100. Park, J., Coddington, L. T. & Dudman, J. T. Basal Ganglia Circuits for Action Specification. Annual Review of Neuroscience vol. 43 485–507 at 10.1146/annurev-neuro-070918-050452 (2020).

101. Yin, H. H. Action, time and the basal ganglia. Philosophical Transactions of the Royal Society B: Biological Sciences vol. 369 at 10.1098/rstb.2012.0473 (2014).

102. Hardcastle, K. et al. Differential kinematic coding in sensorimotor striatum across species-typical and learned behaviors reflects a difference in control. bioRxiv 2023.10.13.562282 (2023) doi:10.1101/2023.10.13.562282.

103. Oldenburg, I. A. & Sabatini, B. L. Antagonistic but Not Symmetric Regulation of Primary Motor Cortex by Basal Ganglia Direct and Indirect Pathways. Neuron 86, 1174–1181 (2015).

104. Hoover, J. E. & Strick, P. L. The organization of cerebellar and basal ganglia outputs to primary motor cortex as revealed by retrograde transneuronal transport of herpes simplex virus type 1. J. Neurosci. 19, 1446–1463 (1999).

105. Alexander, G. E., DeLong, M. R. & Strick, P. L. Parallel Organization of Functionally Segregated Circuits Linking Basal Ganglia and Cortex. Annu. Rev. Neurosci. 9, 357–381 (1986).

106. Aoki, S. et al. An open cortico-basal ganglia loop allows limbic control over motor output via the nigrothalamic pathway. Elife 8, (2019).

107. Mannella, F. & Baldassarre, G. Selection of cortical dynamics for motor behaviour by the basal ganglia. Biol. Cybern. 109, 575–595 (2015).

108. Logiaco, L., Abbott, L. F. & Escola, S. Thalamic control of cortical dynamics in a model of flexible motor sequencing. Cell Rep. 35, 109090 (2021).

109. Rikhye, R. V., Gilra, A. & Halassa, M. M. Thalamic regulation of switching between cortical representations enables cognitive flexibility. Nat. Neurosci. 21, 1753–1763 (2018).

110. Schmitt, L. I. et al. Thalamic amplification of cortical connectivity sustains attentional control. Nature 545, 219–223 (2017).

111. Pearson, K. G., Acharya, H. & Fouad, K. A new electrode configuration for recording electromyographic activity in behaving mice. J. Neurosci. Methods 148, 36–42 (2005).

112. Pachitariu, M., Steinmetz, N., Kadir, S., Carandini, M. & D. H. K. Kilosort: realtime spike-sorting for extracellular electrophysiology with hundreds of channels. bioRxiv 061481 (2016) doi:10.1101/061481.

113. Gage, G. J., Stoetzner, C. R., Wiltschko, A. B. & Berke, J. D. Selective Activation of Striatal Fast-Spiking Interneurons during Choice Execution. Neuron 67, 466– 479 (2010).

114. McInnes, L., Healy, J. & Melville, J. UMAP: Uniform Manifold Approximation and Projection for Dimension Reduction. (2018).

115. Altan, E., Solla, S. A., Miller, L. E. & Perreault, E. J. Estimating the dimensionality of the manifold underlying multi-electrode neural recordings. PLOS Comput. Biol. 17, e1008591 (2021).

116. Pohlmeyer, E. a, Solla, S. a, Perreault, E. J. & Miller, L. E. Prediction of upper limb muscle activity from motor cortical discharge during reaching. J. Neural Eng. 4, 369–379 (2007).

117. Naufel, S., Glaser, J. I., Kording, K. P., Perreault, E. J. & Miller, L. E. A muscle- activity-dependent gain between motor cortex and emg. J. Neurophysiol. 121, 61– 73 (2019).

118. Ehrlich, D. B., Stone, J. T., Brandfonbrener, D., Atanasov, A. & Murray, J. D. PsychRNN: An accessible and flexible python package for training recurrent neural network models on cognitive tasks. eNeuro 8, 1–11 (2021).

